# A multi-phase multi-objective genome-scale model shows diverse redox balance strategies in yeasts

**DOI:** 10.1101/2021.02.11.430755

**Authors:** David Henriques, Romain Minebois, Sebastian Mendoza, Laura G. Macías, Roberto Pérez-Torrado, Eladio Barrio, Bas Teusink, Amparo Querol, Eva Balsa-Canto

**Affiliations:** (Bio)process Engineering Group, IIM-CSIC, Vigo, Spain; Department of Biotechnology, Instituto de Agroquímica y Tecnología de los Alimentos (IATA-CSIC), Paterna, 46980, Spain; Systems Biology Laboratory, VU University, Amsterdam, 1081 HZ, The Netherlands; Department of Genetics, University of Valencia, Burjassot, 46100, Spain

**Keywords:** Batch fermentation, cryotolerant species, dynamic genome-scale models, redox balance, *Saccharomyces* species

## Abstract

Yeasts constitute over 1500 species with great potential for biotechnology. Still, the yeast *Saccharomyces cerevisiae* dominates industrial applications and many alternative physiological capabilities of lesser-known yeasts are not being fully exploited. While comparative genomics receives substantial attention, little is known about yeasts’ metabolic specificity in batch cultures. Here we propose a multi-phase multi-objective dynamic genome-scale model of yeast batch cultures that describes the uptake of carbon and nitrogen sources and the production of primary and secondary metabolites. The model integrates a specific metabolic reconstruction, based on the consensus Yeast8, and a kinetic model describing the time-varying culture environment. Besides, we proposed a multi-phase multi-objective flux balance analysis to compute the dynamics of intracellular fluxes. We then compared the metabolism of *S. cerevisiae* and *S. uvarum* strains in wine fermentation. The model successfully explained the experimental data and brought novel insights into how cryotolerant strains achieve redox balance. The proposed modeling captures the dynamics of metabolism throughout the batch and offers a systematic approach to prospect or engineer novel yeast cell factories.

## 1 INTRODUCTION

Yeasts are among the oldest and most frequently used microorganisms in biotechnology. Approximately 1500 species of yeasts have been described so far. Still, the yeast *Saccharomyces cerevisiae* is the dominant yeast for biotechnology applications. It has been used to produce fermented foods and beverages for millennia and in last years, to produce biofuel (van Zyl et al., 2007), glycerol (Klein et al., 2017), biopharmaceutical proteins (Wang et al., 2017), or secondary metabolites, such as aroma or bioflavours (van Wyk et al., 2018). While many research efforts focus on engineering *S. cerevisiae* strains for particular applications (Nielsen and Keasling (2016); Lian et al. (2018); Liu et al. (2019b); Guo et al. (2020), to name a few), many alternative physiological capabilities of lesser-known yeasts are not being fully exploited (Steensels and Verstrepen, 2014; Hittinger et al., 2015).

Today, genome sequences are publicly available for hundreds of experimentally described yeast species (Dujon and Louis, 2017). These genomic resources have been used to increase our understanding of how different yeasts adapt to natural and anthropogenic environments (Riley et al., 2016). However, non-conventional yeasts provide an unexplored and mostly untapped resource of alternative metabolic routes for substrate utilization and product formation as well as tolerances to specific stressors (Hittinger et al., 2015). Therefore, understanding how the diversity of species and strains use metabolism to grow and produce compounds of industrial and biotechnological interest is essential. Given the complexity of this endeavor, a mathematical modeling approach becomes indispensable.

Genome-scale models (GEMs) can contextualize high-throughput data and predict genotype-environment–phenotype relationships (Oberhardt et al., 2009; Sánchez et al., 2017). While GEMs have been widely used for the analysis of metabolism and the metabolic engineering of *S. cerevisiae* strains (see the recent review by Lopes and Rocha (2017)) in continuous (steady-state) fermentations, their use to predict batch fermentation is still scarce. Nevertheless, many yeast-based processes operate in batch mode.

In batch operation, cell cultures follow a growth curve typically divided into five phases: lag-phase, exponential growth, growth under nutrient limitation, stationary phase and cellular decay. Available dynamic genome-scale models of yeast metabolism focus on the exponential phase and explain reasonably well the measured dynamics of biomass growth, carbon sources uptake and the production of relevant primary metabolites (Hjersted et al., 2007a; Vargas et al., 2011; Sánchez et al., 2014; Saitua et al., 2017a). The development of an integrative modeling approach that describes the five phases of batch processes, considers carbon and nitrogen metabolism throughout time, and explains secondary metabolites’ production is still required.

Here we propose a multi-phase multi-objective dynamic genome-scale model of yeast batch fermentation. The model describes the dynamics of the consumption of carbon and nitrogen (organic and inorganic) sources and the production of primary and secondary metabolites. To model these complex dynamics, we integrated a specific metabolic reconstruction, based on an extension of the current consensus genome-scale model of *S. cerevisiae* (Yeast8, Lu et al. (2019)), into a dynamic kinetic model to account for the time-varying culture environment. We used extracellular substrate concentrations to compute time-varying substrate uptake rates through kinetic expressions. The dynamics of extracellular products was described using kinetic models and used to constrain the problem further. We proposed a multi-phase multi-objective implementation of a parsimonious flux balance analysis (pFBA, Lewis et al. (2010)) to compute the dynamics of the intracellular fluxes.

To illustrate the model’s performance and the novel biological insights that can be gained from its use, we considered a relevant biotechnology process: winemaking. Currently, winemakers can choose among hundreds of *S. cerevisiae* starters (Pretorius, 2000; Matallana and Aranda, 2017; Alonso-del Real et al., 2017). However, to achieve a wider variety of products, other yeasts species are being explored. For example, recent studies revealed that *S. uvarum* strains show interesting physiological properties relevant to wine producers (Querol et al., 2018). *S. uvarum* produces more glycerol and less ethanol than *S. cervisiae* wine strains and the aroma profiles are also different (Masneuf-Pomarède et al., 2010; Gamero et al., 2013; Goold et al., 2017; Alonso-del Real et al., 2017; Varela et al., 2017). In addition, *S. uvarum* is more cryotolerant than *S. cerevisiae*, with an optimal growth temperature around 25*° C* (Belloch et al., 2008; Salvadó et al., 2011), a valuable phenotype in low temperature wine fermentation, which reduces aroma loss. While physiology has been explored for several strains, little is known about the metabolic specificity and flux dynamics of *S. uvarum* under oenological conditions. Recent metabolomic studies have pointed out a higher activity of some relevant pathways in representative strains of *S. uvarum* both in chemostat and batch fermentations (López-Malo et al., 2013; Minebois et al., 2020a). In this work, we applied the proposed model to investigate the origin of such phenotypic divergence.

The model explained the experimental data for all strains successfully and brought novel insights into how these achieve redox balance during wine fermentation. In particular, we hypothesize that cryotolerant yeast strains can use the GABA shunt as an alternative NADPH source and store reductive power (necessary to subdue the oxidative stress under cold conditions) in lipids or other polymers. Additionally, intracellular flux predictions are compatible with recent experimental work showing that most carbon skeletons used to form higher alcohols (e.g., isoamyl alcohol, isobutanol and 2-phenylethanol) are synthesized *de novo*.

Multi-phase and multi-objective dynamic genome-scale models can derive a comprehensive picture of the main steps occurring inside the cell during batch cultures. It is important to stress that the proposed modeling has the potential to capture the dynamics of a wide range of batch processes, in different media and started with various species of yeast, offering a systematic approach to prospect and engineer novel yeast cell factories.

## 2 RESULTS

We derived a multi-phase and multi-objective dynamic genome-scale model of batch fermentation, which accounts for the dynamics of the consumption of carbon and nitrogen (organic and inorganic) sources and secondary metabolites formation. The modeling approach required various refinements to succeed:

i. A novel metabolic reconstruction incorporating missing metabolites and reactions in the Yeast8 consensus reconstruction (see Figure 1.A).
ii. A bootstrap parameter estimation approach to integrate the model with HPLC experimental data to obtain accurate simulations of the secondary metabolism associated with higher alcohols, esters, and carboxylic acids of interest in the biotechnology industry (see Figure 1.A).
iii. A multi-phase, multi-objective dynamic flux balance analysis framework. Different cellular objectives characterized the various phases of the process (lag, exponential growth, growth under nitrogen limitation, stationary, and decay; see Figure 1.B).
iv. A model of protein turnover, required to account for nitrogen homeostasis in the stationary phase (see Figure 1.B).
v. A dynamic biomass equation to better characterize nitrogen assimilation. (see Figures 1.C).

**FIGURE 1.**
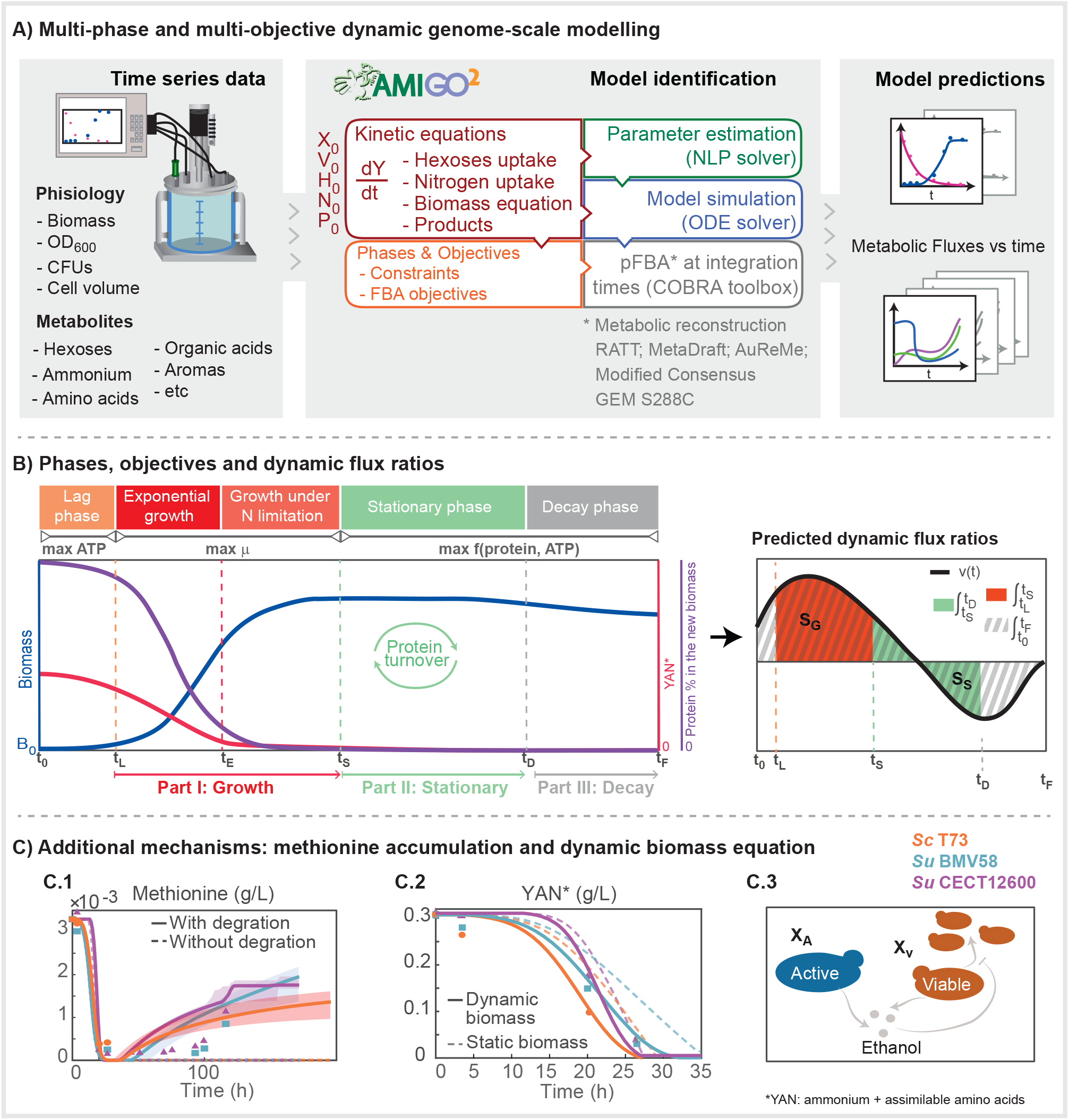
Details on the implementation of the multi-phase and multi-objective dynamic genome-scale model to simulate batch fermentation. A) Implementation, model formulation and solution approach; B) Multi-phase and multi-objective dynamic FBA approach and methodology to compute dynamic flux rates; C) Additional mechanisms included in the model C1) comparison of the model prediction and the experimental dynamics of methionine with and without the methionine degradation pathway, C2) comparison of the performance of the prediction of YAN consumption with static and dynamic biomass equations, C3) schematic view of active vs. viable biomass in the model.

We applied the model to explore the differences in the dynamics of metabolism of *S. cerevisiae* and *S. uvarum* strains in a rich medium (grape juice) fermentation. The final model describes the dynamics of extracellular metabolites successfully with good reproducibility of the experimental data for 46 measured states, including biomass, colony forming units (CFUs), sugars, amino acids and inorganic nitrogen, alcohols and higher alcohols, esters and carboxylic acids. The mean *R*^2^ over the measured states was ⪆ 0.9 for all strains.

We compared the predicted intracellular flux dynamics for the illustrative strains using dynamic flux ratios as computed for two different parts of the fermentation: I) growth - including exponential growth and growth under nitrogen limitation-from *t*_*L*_ to *t*_*S*_ ; II) stationary, from *t*_*S*_ to *t*_*D*_ (see Figure 1.B). The overall dynamic flux ratios from *t_O_* to *t*_*F*_ were also obtained (see details in materials and methods). The major differences between strains appeared in the stationary part, revealing that *S. uvarum* and *S. cerevisiae* strains use different redox balance strategies during the stationary phase of the fermentation:

i. *S. uvarum* generated a large fraction of required NADPH through the GABA shunt;
ii. *S. uvarum* consumed acetate and directed some of this carbon towards mevalonate;
iii. the vast majority of carbon skeletons used to form the higher alcohols isoamyl alcohol, isobutanol and 2-phenyl ethanol were synthesized *de novo*.

### 2.1 Multi-phase multi-objective dynamic genome-scale model

#### 2.1.1 The novel metabolic reconstruction

The Yeast8 model required metabolic refinements to explain secondary metabolism. In total, 38 metabolites and 50 reactions were added to the Yeast8 consensus genome-scale reconstruction of *S. cerevisiae* S288C (v.8.3.1) (Lu et al., 2019) (the list of additions is provided in Table EV1). Furthermore, comprehensive metabolic annotations, such as BO terms and MetaNetX identifiers, were added to the new metabolites and reactions.

Among the metabolites added, 13 aroma compounds were included based on literature data (López-Malo et al., 2013), namely, methionol, ethyl-hexanoate, ethyl-octanoate, 1-hexanol, 1-octanol, 4-tyrosol, hexanal, hexyl-acetate, octyl-acetate, benzyl-acetate, 4-hydroxyphenylacetal-dehyde and 3-methylsulfanylpropanal. Table EV2 presents the chemical family (alcohol, ester or aldehyde) and several aroma descriptors for each of the added compounds. We found that prior genome-scale reconstructions lacked methionol - widely distributed aroma constituent of foods and beverages and the third most relevant aliphatic higher alcohol in wine (De-La-Fuente-Blanco et al., 2016) - and tyrosol - a phenolic higher alcohol known to have antioxidant and anti-inflammatory effects also relevant in wine (Lambrechts and Pretorius, 2000). In the cases of methionol and 4-tyrosol, the lack of these compounds impedes simulated growth on methionine and tyrosine as sole nitrogen sources, which is known to be possible for several *S. cerevisiae* strains and in particular for strain S288C (Cooper, 1982).

The final consensus model was used as a template to reconstruct genome-scale models for wine strains *S. cerevisiae* T73 and *S. uvarum* BMV58 and a strain *S. uvarum* CECT12600 found in non-fermentation environments (strains names will be abbreviated as follows ScT73, SuBMV58 and SuCECT12600 from now on). MetaDraft, AuReMe, and the results from the orthology analysis were used to create the strain-specific models. The three models were almost identical to Yeast8. The models for ScT73, SuBMV58 and SuCECT12600 had 2, 3 and 2 reactions that were not in Yeast8, respectively. A detailed description of the steps followed in the reconstruction and subsequent analyses can be found in the Appendix.

#### 2.1.2 The multi-phase and multi-objective dynamic flux balance analysis

Our results showed that batch fermentation modeling should be divided into five phases in which cellular objectives need to be modified: lag phase, exponential growth, growth under nitrogen limitation, stationary, and decay. The process starts at *t*_0_ = 0 and ends at *t*_*F*_. To define intervals for each phase, we introduced four parameters: *t*_*L*_, *t*_*E*_, *t*_*S*_ and *t*_*D*_ (see Figures 1.A and 1.B for a description of the modeling approach, including the phases and objectives designed in this work).

Once inoculated, cells encounter new nutrients and undergo a temporary period of non-replication, the lag-phase [0-*t*_*L*_], during which we assumed that ATP production is maximized. The exponential growth phase covers only the first hours until nitrogen exhaustion. In this phase [*t*_*L*_−*t*_*E*_], cells maximize growth. The exponential growth phase is followed by growth under nitrogen limitation [*t*_*E*_−*t*_*S*_]; a phase characterized mostly by carbohydrate accumulation. Thenceforward, a substantial fraction of the sugar is consumed during the stationary [*t*_*S*_−*t*_*D*_] and decay [*t*_*D*_−*t*_*F*_] phases by quiescent cells, which adjust their metabolism to cope with environmental fluctuations. In the latter two phases, we assumed cells maximize both ATP and protein production (see further details later in this section). Additionally, depending on the fermentation phase, different constraints were active or inactive.

The general formulation of the FBA problem reads as follows:

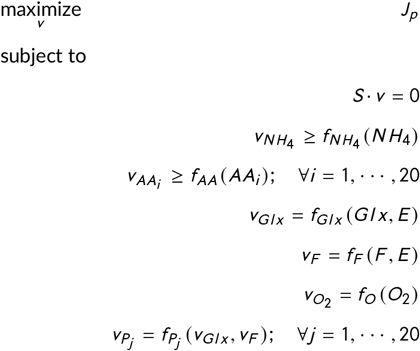

where *J*_*p*_ is the cost functional to be maximized in each phase *p*, *S* is the stoichiometric matrix, *v* is the vector of fluxes in *mmol*/(*g DW h*), *v*_*Gl x*_ and *v*_*F*_ are the fluxes of hexoses (glucose and fructose, respectively), 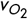 is the flux of *O*_2_ present only at the beginning of the fermentation, 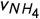 is the flux of ammonium, *v*_*AAi*_ is the exchange rate of the amino acid *i* (with *i* covering all 20 amino acids), *v*_*P*_ are the constraints associated with the 20 fermentation by-products considered here. *Gl x*, *F*, *N H*_4_, *AA*_*i*_, *P*_*j*_ correspond to the concentrations of glucose, fructose, ammonium, amino acids and products, all expressed in (*mmol* /*L*). Expanded View Table EV3 provides a detailed overview of the constraints and the corresponding functional costs at the different phases of the fermentation.

#### 2.1.3 Modelling the dynamics of extracellular metabolites

The uptake of hexoses (glucose and fructose) is modelled using Michaelis-Menten (MM) type kinetics with competitive ethanol inhibition as follows (Hjersted et al., 2007b):

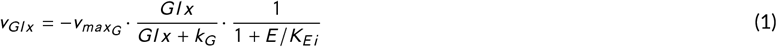

where 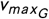 is the maximum rate achieved, *k*_*G*_ is the Michaelis-Menten constant, *K*_*Ei*_ defines the strength of the ethanol inhibitory effect and *E* is the ethanol concentration (mmol/L). A similar expression *v*_*F*_ exists for fructose (*F*). Additionally, in our illustrative examples, the grape juice was supplemented with sucrose; thus, we included a mass action type expression describing its hydrolysis into glucose and fructose. The kinetics of this reaction was controlled by the parameter *k*_*hydro*_.

During the lag-phase our data shows the presence of dissolved oxygen (see Expanded View Figure EV1.B), therefore its consumption was allowed and constrained by a mass action type kinetics:

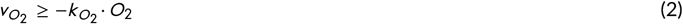

where 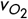 is the oxygen uptake rate, 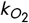 a parameter controlling the oxygen transport rate and *O*_2_ the concentration of oxygen in the media.

We characterized the uptake of different nitrogen sources including ammonium and amino acids. Ammonium kinetics was modeled by a Michaelis-Menten expression:

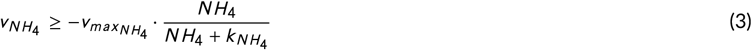

where *N H*_4_ is the extracellular concentration of ammonia (mmol/L), 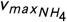 is the maximum uptake rate achieved, 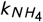 is the Michaelis-Menten constant. To avoid an excessive number of parameters, amino acid transport was modeled following mass action kinetics:

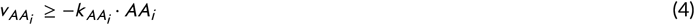

where *AA*_*i*_ is the extracellular concentration of the amino acid (mmol/L) and 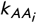 is the associated kinetic parameter.

The extracellular dynamics of most metabolic by-products is governed by:

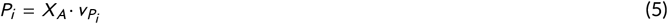

with *X*_*A*_ the active biomass, and the production flux 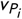 proportional to the amount of transported hexoses:

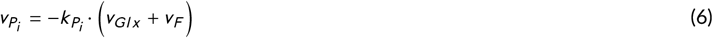

where *P*_*i*_ refers to the excreted product *i* and 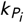 controls the magnitude of the metabolite production.

An exception to this was the formulation of the dynamics of acetate. This metabolite is produced during the exponential growth phase and consumed during the stationary phase following mass action kinetics. The model accounts for the production of alcohols and higher alcohols (ethanol, glycerol, 2,3-butanediol, 2-phenylethanol, isobutanol, isoamyl alcohol, 1-hexanol, benzyl alcohol), carboxylic acids (succinate, malate, acetate and lactate) and esters (ethyl acetate, isobutyl acetate, isoamyl acetate, 2-phenylethyl acetate, ethyl caprate, ethyl caprylate, ethyl caproate and hexyl acetate).

#### 2.1.4 The model of protein turnover

The metabolic model required incorporating a nitrogen source to account for the metabolism of amino acids and their related higher alcohols during the stationary phase. Because all nitrogen sources in the media have been depleted before the stationary phase, we developed a new model of nitrogen homeostasis that considered protein turnover. The proposed model describes the combined use of the Ehrlich and *de novo* synthesis pathways during stationary and decay phases to guarantee optimal adaptation to perturbations in nitrogen homeostasis.

To introduce protein turnover to our model, we simulated the degradation of the existing protein fraction inside biomass (*P r ot*), into a pool of amino acids that subsequently produce new proteins. During stationary and decay phases, the lower bounds on the amino acid uptake are set as:

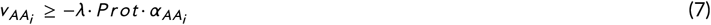

where *λ* is the turnover rate, *P r ot* is the concentration of protein and 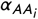 is associated with the stoichiometric coefficient specifying the quantity of a given amino acid in the protein pseudo reaction. Formally, the time-course evolution of protein content is described by the following equation:

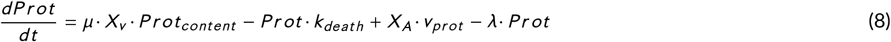

where *μ* corresponds to the growth rate, *P r ot*_*cont ent*_ to the fraction of protein in the newly formed biomass, *k*_*death*_ (*h*^−^1) is a parameter controlling the rate of biomass degradation during decay phase, *X*_*V*_ is the simulated viable biomass (g/L), *X*_*A*_ the simulated active biomass (g/L), *v*_*P r ot*_ the protein production rate (*g* · *g DW*^−1^ · *h*^−1^) and *λ* the protein turnover rate (*h*^−1^).

The degraded proteins (*λ* · *P r ot*) are converted into extracellular amino acids whose concentrations are represented by the following equations:

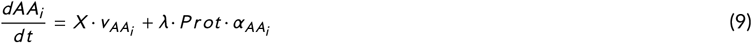

where 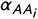 is associated with the stoichiometric coefficient specifying the quantity of a given amino acid in the protein pseudo reaction associated with the flux *v*_*P r ot*_. The rates 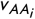 are computed by maximizing protein production (*v*_*P r ot*_) and ATP (*V*_*AT P*_) while solving the FBA problem:

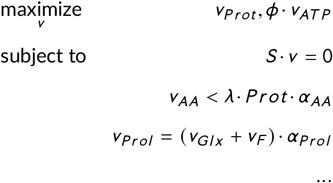

where *φ* is a estimated parameter for each strain, *S* is the stoichiometric matrix, *v* is the vector of fluxes, 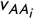 is the exchange rate of amino acid *AA*_*i*_, 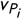 constraints associated with fermentation by-products (see more details in the Methods section) and *v*_*Prol*_ (r_1904) is the amount of excreted proline. The later amino acid accumulates in the extracellular media throughout the fermentation (see Expanded View Figure EV1.A), likely as a consequence of stored arginine consumption in anaerobic conditions (see Crépin et al. (2014)). The extracellular dynamics of proline is described by the equation:

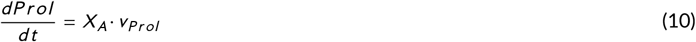

Depending on the kinetic constraints, amino acids can be directly incorporated into proteins or degraded to recover nitrogen for protein production. In line with this, we observed that those amino acids which lacked pathways for their catabolism or elimination of by-products accumulated in the extracellular compartment. For example, the lack of methionol in the Yeast8 model led to the accumulation of methionine during the stationary phase. Figure 1.C1 shows how the model can successfully recover the dynamics of methionine, suggesting the pathway for methionine degradation could be active during the stationary phase.

#### 2.1.5 The dynamic biomass equation

Our initial model with a static biomass equation was not able to reproduce nitrogen assimilation despite parameter and adjustment of protein content (Figure 1.C2). Based on the observations of Varela et al. (2004); Vargas et al. (2011) and our modelling results, we implemented the following dynamic biomass equation:

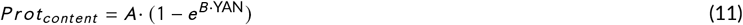

where A and B are estimated parameters and YAN the quantified yeast assimilable nitrogen. YAN accounted for the ammonium and the free amino acids present in the white must, excluding proline, which is not catabolized under anaerobic conditions. Furthermore, we assumed that the mRNA level was proportional to the protein content (*mRN A* = *prot*/*RNA*_*to*_*Protein*_*Ratio*). In this framework, carbohydrates compensate for the variation in protein and mRNA content. Growth-associated ATP maintenance (*GAM*) was also updated to account for the polymerization costs of the different macromolecules (protein, RNA, DNA and carbohydrates):

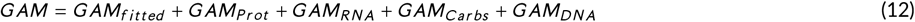

where *GAM*_*f i t t ed*_ is a species or strain-dependent parameter that is estimated and the rest are polymerization costs of the different biomass precursors (as adapted from Lu et al. (2019)). Additionally, to represent the premature end of fermentations during the decay phase (observed in SuCECT12600), we estimated the non–growth-associated maintenance (NGAM). The former formulation, coupled to parameter estimation, allowed us to predict nitrogen consumption dynamics accurately, as shown in Figure 1.C2.

In addition, we discriminated between active and viable cell mass to capture the dynamics of CFUs and biomass (Figure 1.C3). Here, we define active cells as those that are able to ferment and viable cells as those that are able to divide and ferment. The dynamics of active cell mass is represented by the equation:

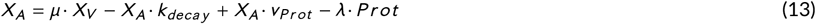

where *X*_*A*_ is the active cell mass (g/L), *μ* is the growth rate computed with pFBA, *k*_*decay*_ is the decay rate (only active during decay phase), *v*_*P r ot*_ (r_4047) the exchange flux for protein production (only active during stationary and decay phases) and *λ* is the turnover rate (only active during stationary and decay phases). The behaviour of viable cell mass differed from that of active cell mass by a decline induced by ethanol (Cramer et al., 2002):

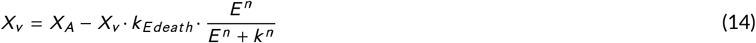

where *X*_*v*_ (g/L) is the growth rate obtained with the constraint based model, *E* is the ethanol concentration (mmol/L) and *k*_*E deat h*_, *n* and *k* are the parameters controlling susceptibility to ethanol.

#### 2.1.6 Goodness-of-fit of the model in the illustrative examples

We integrated the model with experimental data obtained in batch fermentations through a bootstrap parameter estimation approach (Figure 1.A, see Materials and methods). The model described the dynamics of our illustrative examples successfully. The final model consisted of 46 ordinary differential equations, describing the dynamics of the external metabolites, plus the algebraic constraints representing the pseudo-steady state metabolic state at each time step. The model depends on 66 unknown parameters, estimated from the experimental data using a bootstrap approach. The best fit to the data plus the associated uncertainty, as computed by the bootstrap approach, are represented for the biomass, hexoses and few relevant metabolites in Figures 2 and 3. The Expanded View Figure EV1 shows the best fit to the remaining measured metabolites. The Expanded View Figure EV2 presents the parameter estimation convergence curves for the different strains and bootstrap realizations.

**FIGURE 2.**
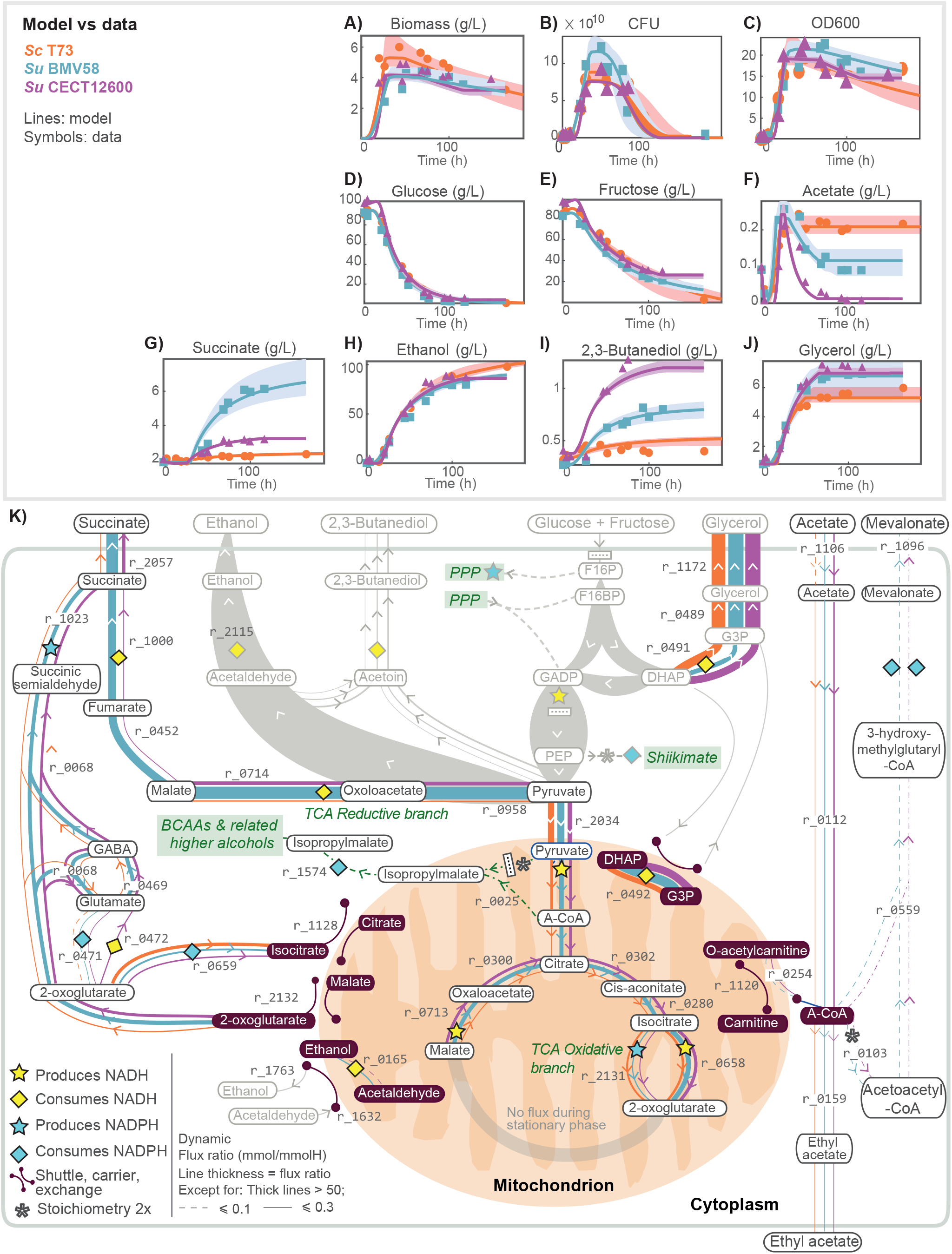
Redox balance in Central Carbon Metabolism. Figures A) to J) show model predictions versus the experimental data for biomass, and extracellular metabolite concentrations associated with glycolysis and central carbon metabolism for the three strains. Figure K) shows the predicted intracellular dynamic flux ratios associated with the central carbon metabolism during the stationary phase. Figure K) illustrates how *S. uvarum* and *S. cerevisiae* strains use different redox balance strategies during the stationary phase of the fermentation. These differences result in the differential production of relevant external metabolites such as ethanol (D), succinate (G), 2,3-butanediol (H), or glycerol (J).

**FIGURE 3.**
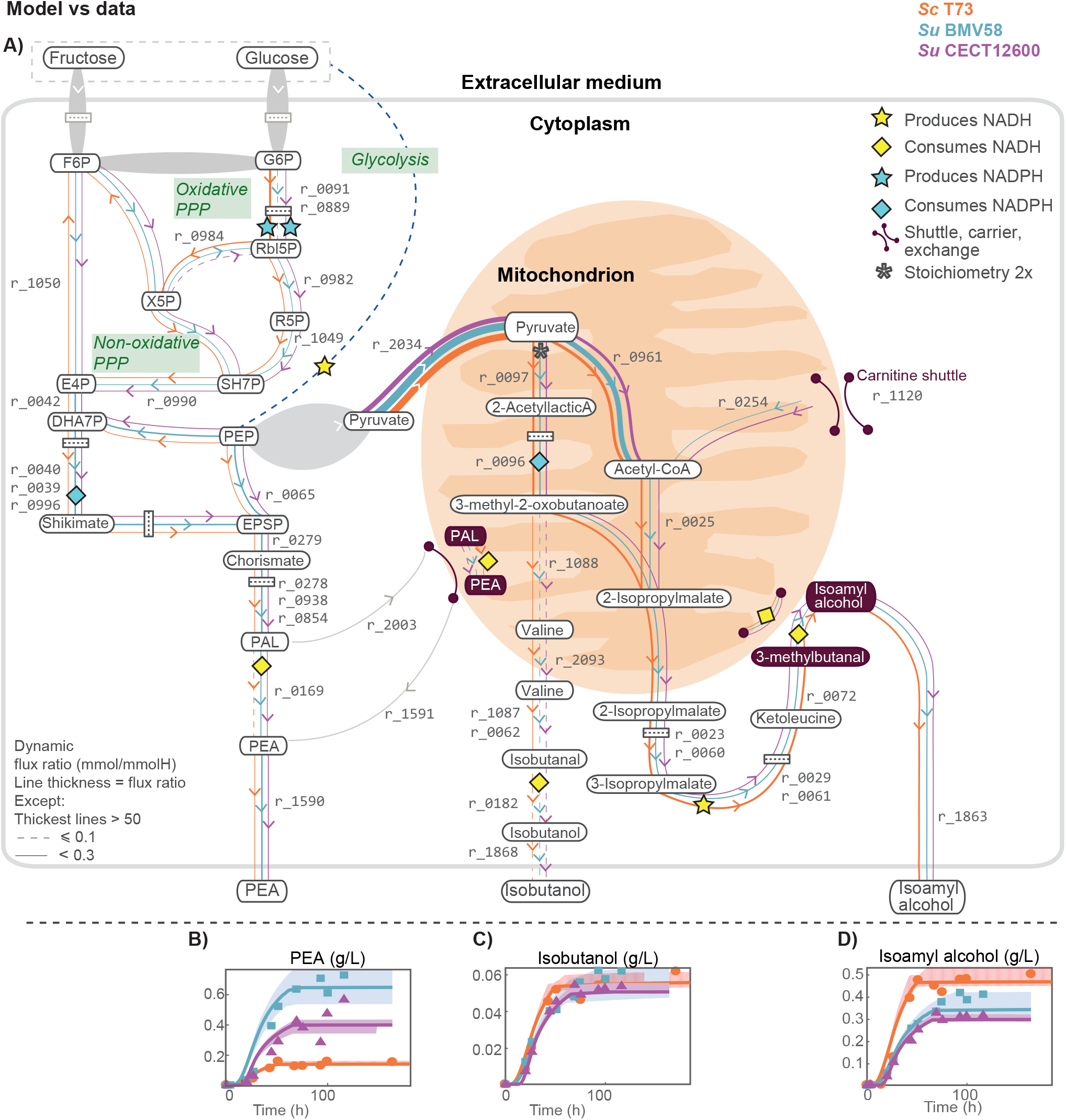
Redox balance in higher alcohol production: A) Fluxes associated with the production of higher alcohols 2-phenylethanol (PEA), isobutanol and isoamyl alcohol during the stationary phase. Figures B) to D) show the model predictions versus the experimental data corresponding to the extracellular concentrations of PEA, isobutanol, and isoamyl alcohol, respectively.

We determined the R-squared measure of goodness of fit (*R*^2^) for each measured variable and each strain-based fermentation. The Expanded View Table EV4 presents the corresponding values. The vast majority of the coefficients were positive with few exceptions (10 out 141), typically associated with low signal-to-noise ratio and high data dispersion observed in the measured variable, e.g., cysteine. The mean *R*^2^ (excluding negative values) was of 0.88 for *S. cerevisiae* T73, 0.92 for *S. uvarum* BMV58 and 0.91 for *S. uvarum* CECT12600; the median *R*^2^ values are above 0.94 for all strains.

The Expanded View Figure EV3 presents the bootstrap parameter distributions showing that uncertainty on the parameter estimates was reasonably low. Its value varied between parameters and species. The mean standard deviation corresponds to a 9.5% for the parameter values estimated for ScT73 (excluding the parameter *N GAM*, whose optimal value was zero). A slightly higher mean standard deviation (12.6%) was obtained for the parameter values estimated for SuBMV58 (excluding the kinetic constant describing benzyl alcohol production whose optimal value was zero), and a significantly lower mean uncertainty (2.5%) was obtained for those parameters corresponding to SuCECT12600. The reasonably low distribution on the parameters resulted in a reasonably low uncertainty associated with the model simulations (as seen in Figures 2,3, and Expanded View Ev1).

Some parameter values vary significantly between strains (Expanded View Table EV5). The comparison between the wine strains ScT73 and SuBMV58, reveals substantial differences - above 100% relative difference, in the growth-associated ATP maintenance; the rate of transport of specific amino acids and hexoses and the production of certain alcohols and acids. *GAM*_*f i t t ed*_ for SuBMV58 is around twice the value for ScT73. The rate of transport of phenylalanine, leucine, alanine and cysteine is between 106% and 778% larger for SuBMV58 than for ScT73. Similarly, hexoses uptake rate is around two times higher at the beginning of fermentation for SuBMV58 than for ScT73.

Regarding products, the most noticeable difference appears in the production rate of succinate and, to a lesser extent, in the production of 2-phenylethanol, 2-phenylethyl acetate, and 2,3-butanediol. The comparison between the wine strains and the natural strain showed that the lag phase is more than three times longer for the natural strain; besides, the non-growth associated maintenance is practically zero for wine strains while it is around 0.8 for the SuCECT12600 strain. The rates of hexoses uptake are pretty similar between *S. uvarum* strains; on the contrary, we found substantial differences in the uptake of various amino acids. In particular, the uptake rate for threonine is 740% higher for SuCECT12600 than for SuBMV58.

### 2.2 Biological insights gained from the model

At the extracellular level, we observed the most striking differences between strains during the stationary phase in the dynamics of acetate and the yield of succinate (see Figure 2) and in the production of higher alcohols (see Figure 3). We used the proposed multi-phase multi-objective dynamic genome-scale model to explore the metabolic pathways associated with these differences. The Expanded view Table EV6 reports the dynamic flux ratios as mmol of produced compound per mmol of consumed hexose x 100 (from now on, mmol/mmolH) for those reactions in which the maximum flux ratio value over the three species is above 0.01 mmol/mmolH.

Our model showed different redox balancing among yeast species in fermentation at the central carbon metabolism level and the production of higher alcohols (interested readers can find a detailed discussion in the Appendix).

The most relevant differences appeared in the use of pyruvate in the mitochondrion. *S. cerevisiae* and *S. uvarum* strains shifted similar quantities of pyruvate towards the mitochondrion (3.2, 3.6, 2.4; r_2034, Figure 2.K). However, its fate appears to differ by species. *S. uvarum* oxidized pyruvate up to 2-oxoglutarate and sent it into the cytoplasm to generate NADPH with the help of the GABA shunt. In contrast *S. cerevisiae* shifted most of this carbon towards the branched-chain amino acid pathway. Accordingly, significant differences were observed in the production of succinate and isoamyl alcohol.

Our results also revealed that *S. uvarum* species consume acetate once nitrogen sources are no longer available. Both species produced acetate during the growth part of the fermentation. However, *S. uvarum* strains consumed part of it during the stationary phase. The model revealed that *S. uvarum* used acetate to produce mevalonate and used the carnitine shuttle system to translocate acetyl units to the mitochondrion. Notably, the model suggested that the production of 2-phenylethanol is also a pivotal contributor in the cellular redox state of *S. uvarum* strains.

#### 2.2.1 The GABA shunt as an NADPH source in cryotolerant species

The model estimated significantly higher dynamic flux ratios of succinate on the overall fermentative process in SuBMV58 (3.59) and SuCET12600 (1.10) than ScT73 (0.35) (Expanded View Table EV6, r_2057). Remarkably, the difference in production between *S. uvarum* strains was noteworthy. Cells produced the most significant fraction of succinate during the stationary and decay phases, with a significantly higher dynamic flux ratio by SuBMV58 and SuCECT1600 (6.08 and 1.66, r_2057) than ScT73 (0.42, r_2057).

During the decay phase, most succinate was produced through the TCA cycle reductive branch in the two species. However, the model predicted that, during the stationary phase, succinate productions was distributed between the GABA shunt (0.42, 2.00 and 1.31 for ScT73, SuBMV58 and SuCECT12600 respectively; r_1023, Figure 2.K) and the reductive branch of TCA (0.00, 4.10 and 0.35 for ScT73, SuBMV58 and SuCECT12600 respectively; r_1000, Figure 2.K). The reductive branch of the TCA cycle contributed almost twice the GABA shunt in succinate formation for SuBMV58. In contrast, the GABA shunt was the primordial source of succinate during the stationary phase in ScT73 and SuCECT12600 strains.

During the stationary phase, the TCA oxidative branch was active up to 2-oxoglutarate (precursor of succinate). The model suggested that, in the absence of oxygen, the NADH produced could be re-oxidized at the level of the mitochondrial shuttles responsible for reducing cytoplasmic dihydroxyacetone phosphate (DHAP) to glycerol 3-phosphate (G3P) (2.69, 4.98 and 3.46, r_0492) and cytoplasmic acetaldehyde to ethanol in the mitochondrion (0, 0.52 and 0.46, r_0165).

The high predicted activity of the GABA shunt in *S. uvarum*, between 3 and 4.6 times higher than *S. cerevisiae* (Figure 2.K, r_0068, r_1023; Expanded View Figure EV4) suggested an important role in the maintenance of the cellular redox state. Incidentally, revisiting data from previous experiments by López-Malo et al. (2013) — who compared fermentations started with *S. uvarum* (SuCECT12600) and *S. cerevisiae* (QA23)— we found a considerable accumulation of GABA (93.59 fold-change) by SuCECT12600 (see supporting information Table EV7).

#### 2.2.2 Intracellular mevalonate as a reducing equivalent in cryotolerant yeast species

The three strains produced acetate during the growth phase and until the entry into the stationary phase. We obtained quite a similar dynamic flux ratio towards acetate production during the growth phase in the three strains (1.079, 1.145, 1.422 in ScT73, SuBMV58 and SuCECT12600, respectively; Expanded View Table EV6,r_1106). Afterward, while extracellular acetate concentration remained constant in ScT73, a decrease was observed in both *S. uvarum* fermentations, indicating acetate consumption. As shown in Figure 2.F, our model successfully described these phenotypes.

*S. uvarum* and *S. cerevisiae* produced similar amounts of ethyl acetate (0.14, 0.17 and 0.19 in ScT73, SuBMV58 and SuCECT12600, respectively; Expanded View Table EV6,r_0159). However, *S. uvarum* strains also used the carnitine shuttle system to transport acetyl-CoA into the mitochondria (0.09 and 0.38 in SuBMV58 and SuCECT12600, respectively; Figure 2.K; Expanded View Table EV6, r_0254). Inside the mitochondria, acetyl-CoA was used to form isopropylmalate (Figure 3.A) - a precursor of leucine and isoamyl alcohol - or in the TCA oxidative branch towards the synthesis of 2-oxoglutarate (Figure 2.K).

In our first simulations, the model determined that cells use a significant part of the consumed acetate to produce succinate through the glyoxylate pathway. However, we could not find literature supporting the production of glyoxylate in the presence of a large glucose concentration. Therefore, we decided to constrain the isocitrate lyase flux to zero. The revised model suggested that during the stationary phase *S. uvarum* strains incorporated the acetate derivative, acetyl-CoA, into mevalonate (0.19 and 0.10 in SuBMV58 and SuCECT12600, respectively; Expanded View Table EV6,r_0559 and r_0103) also consuming NADPH. It should be noted that the pathway from acetyl-CoA to mevalonate is reversible. In addition, mevalonate is a reducing equivalent that can be further metabolized into, for example, ergosterol (r_0127 in our reconstruction) possibly acting as storage of NADPH in cryotolerant species.

#### 2.2.3 The production of higher alcohols contributed to the redox balance

Higher alcohol production was most prominent during the stationary phase for the three strains. Our model predicted that carbon skeletons of isoamyl alcohol, isobutanol and 2-phenylethanol (PEA) were in great part synthesized *de novo* from glycolytic and pentose phosphate pathway intermediates, rather than coming from the catabolism of precursor amino acids (leucine, valine and phenylalanine respectively). Figure 3.A shows the predicted intracellular flux ratios (above 0.01 mmol/mmolH) related to higher alcohols during the stationary phase and their corresponding impact on the redox co-factors balance NADPH/NADP+ and NADH/NAD+. Figures 3.B-3.D correspond to the comparison between model predictions and raw measures of PEA, isobutanol and isoamyl alcohol, respectively. The Expanded View Figure EV4 and Table EV6 present the dynamic flux ratios. Remarkably, the flux ratios corresponding to the degradation of amino acids are well below 0.01. During the stationary phase, *S. uvarum* strains produced more PEA than ScT73 strain per unit of consumed hexoses (0.158, 0.659, 0.383 for ScT73, SuBMV58 and SuCECT12600, respectively; r_1590) while the opposite pattern was observed for isoamyl alcohol (0.736,0.483, 0.394 for ScT73, SuBMV58 and SuCECT12600, respectively; r_1863), although differences were less noticeable. In contrast, the model prediction was quite similar for the three strains for isobutanol, as well as for other higher alcohols such as methionol and tyrosol, which seemed to accumulate in minimal quantities (flux ratio *x* 100 ⪆ 0.001) in response to perturbations in the amino acid pool.

We found that the production of higher alcohols contributed substantially to the redox metabolism related to glycerol accumulation. During the stationary phase, approximately 43% of the glycerol produced by the ScT73 strain was attributable to NADH derived from isoamyl alcohol and PEA. In the cases of SuBMV58 and SuCECT12600 strains, these values dropped to 36% and 27%, respectively (see further details in the Appendix).

As shown in Figure 3.A, our model predicted that during the stationary phase, ScT73 had a larger flux ratio through the oxidative pentose phosphate pathway (PPP, r_0091, r_0889) than *S. uvarum* strains. Fluxes through the shikimate pathway were higher in both *S. uvarum* strains (Figure 3.A, r_0996 and r_0279), an increase associated with the higher production of PEA observed in *S. uvarum* strains. Interestingly, while ScT73 seemed to redirect part of the oxidative PPP flux towards glycolysis, *S. uvarum* simulations reflected the inverse pattern (r_0984, Figure 3.A), with glycolytic flux being shifted towards the non-oxidative PPP.

Pyruvate in the mitochondrion showed two different fates (Figure 3.A): it can be converted into acetyl-CoA (r_0961) and used to produce 2-acetyllactic acid (r_0097). Noticeably, *S. uvarum* strains also contributed to acetyl-CoA using the carnitine shuttle (r_0254). 2-acetyllactic acid can further be converted to 3-methyl-2-oxobutanoate, consuming one NADPH (r_0096), which also showed two different fates. On the one hand, it can be directed toward 2-isopropylmalate (r_0025), leading to the formation of cytosolic ketoleucine, the precursor of isoamyl alcohol (r_0072, r_0179). On the other hand, 3-methyl-2-oxobutanoate can be used as the precursor of valine and for the synthesis of isobutanol via the Ehrlich pathway (r_1087, r_0062, r_0182). Similar amounts of isobutanol were exported by all three strains (0.1, 0.084, 0.078; r_1868).

## 3 DISCUSSION

Many recent studies have discussed the possibility of finding yeasts with novel phenotypes for optimized biotechnological applications. Strategies for finding such yeasts include bioprospecting for novel strains and species (Steensels and Verstrepen, 2014; Pérez-Torrado et al., 2018), hybridization and evolution of known yeast strains or species (Peris et al., 2017) and the synthetic design of novel properties (Deparis et al., 2017; Löbs et al., 2017; Mans et al., 2018). With the development of omics analysis and genome-scale modeling tools, it is possible to characterize and even design phenotypes of microorganisms to serve as efficient cell factories (Borodina and Nielsen, 2014).

Prior studies have considered batch fermentation modeling using available yeast metabolic reconstructions and dynamic FBA implementations (Hjersted et al., 2007a; Vargas et al., 2011; Vázquez-Lima et al., 2014; Saitua et al., 2017b). However, these studies focused on the exponential growth phase, thus not explaining secondary metabolism. Their static nature would also prevent the accurate description of the sequential nature of amino acid consumption. Therefore the final models were not able to fully capture fermentation dynamics nor relevant secondary metabolites.

This study aimed to develop a dynamic genome-scale model to investigate the dynamics of yeasts primary and secondary metabolism in batch cultures. The first question in this research was how to describe all phases in the batch process: lag, exponential growth, limited nitrogen growth, stationary and decay. To generate biological hypotheses for the modeling and to test its predictive nature, we considered the dynamics of metabolism of *S. cereivisiae* and *S. uvarum* strains in a rich medium (grape juice) fermentation.

The current study found that the model requires four new features to describe batch fermentation metabolism successfully:

i. **A specific metabolic reconstruction** that incorporates several missing metabolites and reactions in the Yeast8 consensus model. A similar curation process was applied to the iMM904 reconstruction (Scott et al., 2020). The integration with the corresponding HPLC experimental data allows obtaining more accurate simulations of the secondary metabolism associated with higher alcohols, esters, and carboxylic acids.
ii. **A multi-phase multi-objective dynamic FBA scheme** to distinguish the five phases of the process (lag-phase, exponential growth, growth under nitrogen limitation, stationary, and decay). Each phase is characterized by a different cellular objective: maximum growth, maximum ATP production, maximum protein production. Remarkably, while previous works focused on ATP consumption to explain the metabolism after depletion of the limiting nutrient (Raghunathan et al., 2006; Pizarro et al., 2007; Vargas et al., 2011), we incorporated the production of protein as a cellular objective (in conjunction with protein degradation) with accurate results.
iii. **A model of protein turnover** to describe the uptake of amino acids and inorganic nitrogen. Protein turnover in the stationary phase explains nitrogen homeostasis, in which the removal of any amino acid corresponds to a perturbation. To the best of our knowledge, this is the first dynamic genome-scale metabolic model describing nitrogen homeostasis during the stationary phase.
iv. **A dynamic biomass equation** that considers the dynamics of inorganic and organic nitrogen sources in the medium over time. Previous works have shown the dynamics of biomass composition Schulze et al. (1996) or Varela et al. (2004), and Dikicioglu et al. (2015) pointed out the relevance of detailing its composition in a context-specific manner.

The second question in this research was to decipher the differences in the metabolism of three strains of two different species, *S. cerevisiae* and *S. uvarum*, in wine fermentation. In a previous study, Minebois et al. (2020b) hypothesized that *S. cerevisiae* and *S. uvarum* species may have different redox homeostasis strategies. In the present work, we confirmed this hypothesis. Notably, the model brought novel insights into the different redox balance strategies used by *S. cerevisiae* and cryotolerant *S. uvarum* strains.

Our flux predictions, together with previous findings, suggest alternative pathways for cryotolerant species to produce succinate and consume acetate. In principle, yeasts might form succinate via four main pathways, all based on the reactions of the TCA cycle (Coulter et al., 2004): the TCA cycle, the GABA shunt, the glyoxylate cycle and the methylcitric acid cycle, depending on the environmental conditions and strain. Our model revealed that, at the early stages of fermentation, when there was a small amount of oxygen present in the medium, the TCA pathway operated as a cycle. However, once oxygen was depleted, the TCA pathway did not longer work as a cycle but in a branched manner. In that scenario, the model predicted that on the overall fermentative process, cells produced most succinate via the TCA reductive branch, in agreement with Camarasa et al. (2003). However, our results also suggest an important role of the TCA oxidative branch until 2-oxoglutarate for the *S. uvarum* strains during the stationary phase. This result is consistent with the recent intracellular data obtained by Minebois et al. (2020a) who observed a noticeable intracellular accumulation of this TCA intermediary in the stationary phase of SuBMV58.

One somewhat unexpected finding of the model was the extent to which the GABA shunt would contribute to succinate formation during the stationary phase; an effect particularly evident in the case of *S. uvarum* strains. The role of this pathway is not fully understood in yeast (Mara et al., 2018; Cao et al., 2013; Bach et al., 2009). In a previous study, Bach et al. (2009) observed that glutamate decarboxylase (*GAD1*) was poorly expressed in *S. cerevisiae* when succinate was produced and that the GABA shunt played a minor role in redox metabolism. However, Coleman et al. (2001) showed that glutamate decarboxylase (*GAD1*) is required for oxidative stress tolerance in *S. cerevisiae*. Later, Cao et al. (2013) showed that *GAD1* confers resistance to heat stress by reducing reactive oxygen species while hypothesizing that this effect could be related to NADPH production. Additionally, *GAD1* was found to be up-regulated during the stationary phase, under nitrogen starvation (Mara et al., 2018; Bach et al., 2009; Rossignol et al., 2003); and López-Malo et al. (2013) observed high intracellular GABA levels in cryotolerant species *S. uvarum*.

Recently, Liu et al. (2019a) found clear indication that the GABA shunt may be involved in supplying NADPH for lipid synthesis in the oleaginous yeast *Yarrowia lipolytica*. Also, Bach et al. (2009) presented evidence that *S. cerevisiae* can degrade GABA into succinate or *γ*-Hydroxybutyric acid (GHB) and that GHB was used to form the polymer polyhydroxy-butyrate (PHB). Strikingly, Bach et al. (2009) found that *S. cerevisiae* formed PHB by a mixture of GHB (GABA catabolism) and 3-hydroxybutyrate-CoA (derived from acetyl-CoA), with a ratio that was dependent on GABA supplementation. Also noteworthy is that PHBs are synthesized by numerous bacteria as carbon and energy storage compounds (Możejko-Ciesielska and Kiewisz, 2016), acting as dynamic reservoirs of carbon and reducing equivalents. In these species, PHBs play a critical role in central metabolism and typically accumulates under conditions of nutritional imbalance (López et al., 2015); PHBs are also strongly associated with bacterial cold tolerance (Müller-Santos et al., 2020) suggesting a common function in yeast.

Another important finding is that *S. uvarum* strains consume acetate once nitrogen sources are depleted. Previous studies (Giudici et al., 1995; Bertolini et al., 1996) reported that *S. uvarum* produces less acetate than *S. cerevisiae*. Actually, by following the time-course of the fermentation, we observed that both species appear to produce similar amounts of acetate during the growth phase, but *S. uvarum* strains consume all (SuCECT12600) or a significant fraction (SuBMV58) during the stationary phase, coinciding with the extracellular accumulation of succinate. This finding was also recently reported by Kelly et al. (2020), who showed that a *S. uvarum* yeast isolate can metabolize acetate to significantly lower acetic acid, ethyl acetate, and acetaldehyde in wine. According to the modeling constraints, the most parsimonious explanation for this observation would have been an operative glyoxylate cycle. However, based on the repression by glucose of the key enzymes of the glyoxylate cycle (i.e. *ICL1* and *MLS1*) and previous intracellular data (López-Malo et al., 2013), we decided to block this cycle and explored an alternative hypothesis. In this new scenario, the model predicted that some of the acetate carbon was directed towards mevalonate, which is in line with recent experimental work by (Minebois et al., 2020a). Following the model proposed by Bach et al. (2009), a route for acetyl-CoA incorporation into PHB polyester through 3-hydroxybutyrate-CoA seems plausible. However, this hypothesis is not taken into account by the genome-scale reconstructions.

The fact that López-Malo et al. (2013) found high intracellular GABA and GHB in cryotolerant strains grown in synthetic must (without GABA) at low-temperature, and the flux predicted in the present work, suggest that *S. uvarum* stores lipids or polyesters (i.e., PHBs) as reducing equivalents to withstand oxidative stress induced by low temperatures, a conclusion that requires further investigation. Approaches such as the one proposed by Paget et al. (2014) might shed further light on the thermodynamics of the different pathways involved in the cryotolerant phenotype.

Our model also predicted that the carbon skeletons of higher alcohols (e.g., isobutanol and isoamyl alcohol) were mainly synthesized *de novo* rather than proceeding from the incorporation and catabolism of precursor amino-acids (e.g., leucine and valine). This result agrees with the findings of Crépin et al. (2017) who explored the fate of the carbon backbones of aroma-related exogenous amino acids using 13C isotopic tracer experiments. Similarly, our results indicate that 2-phenylethanol was mostly synthesized *de novo* and not derived from exogenous amino acid catabolism through the Ehrlich pathway. We hypothesize that a positive contribution in glycerol content may also explain why the production of 2-phenylethanol and isoamyl alcohol is a conserved evolutionary trait in yeasts.

Under the experimental conditions, isoamyl alcohol and 2-phenylethanol significantly impacted redox metabolism, mainly when biosynthetic requirements were low (late growth phase and stationary phase). Despite that the NADH requirements of 2-phenylethanol are four times smaller than those of isoamyl alcohol, the former also impacted the NADP+/NADPH ratio. The PPP branch followed to produce 2-phenylethanol precursor erythrose-4 phosphate might result in NADPH or NADP+ accumulation. Our model predicted that during the stationary phase, 2-phenylethanol produced through the chorismate synthesis pathway (downstream of the non-oxidative PPP), provided some of the excess NADP+ required to transform succinic semialdehyde into succinate. The present model (along with the provided code) can be used to assess the redox contribution of this and alternative explanations.

Interestingly, the model also predicted that most other higher alcohols (tyrosol, methionol, etc.) accumulated in small amounts due to perturbations in the amino acid pool. These results corroborate the ideas of Stevens (1961); Shopska et al. (2019) who suggested that higher alcohols in beer were by-products of amino-acid and protein synthesis and that the two schemes of production (Ehrlich and *de novo* synthesis) were not in contradiction but were two extremes of a common mechanism. Incidentally, recent research by Yuan et al. (2017) showed that the assembled leucine biosynthetic pathway coupled with the Ehrlich degradation pathway resulted in high-level production of isoamyl alcohol.

Summing up, this work addressed two fundamental questions. The first relates to the possibility of describing the dynamics of the primary and secondary metabolism of yeasts in batch cultures; the second, to the biological insights that can be gained from the use of such a model. The present model (along with the provided code) can be used to simulate yeast metabolism in batch culture; the only requirement would be to update the metabolic reconstruction if required for the specific yeast species. The model can also be used to explore and engineer novel metabolic pathways towards specific bioproducts. The fact that model predictions are consistent with numerous previous findings led us to conclude that other novel results, such as the role of the GABA shunt and the production of reducing equivalents in the metabolism of cryotolerant species may be plausible routes worth exploring.

## 4 MATERIALS AND METHODS

### 4.1 Yeast strains

In this study, three yeast strains belonging to *S. cerevisiae* and *S. uvarum* species were used: the commercial strain, T73 (Lalvin T73 from Lallemand Montreal, Canada), originally isolated from wine in Alicante, Spain (Querol et al., 1992) was selected as our wine *S. cerevisiae* (ScT73) representative; the commercial strain BMV58 (SuBMV58, Velluto BMV58 from Lallemand Montreal, Canada), originally isolated from wine in Utiel-Requena (Spain) and the non-commercial CECT12600 strain, isolated from a non-fermentative environment (SuCECT12600, Alicante, Spain).

### 4.2 Wine fermentation experiments

Fermentation assays were performed with must obtained from the Merseguera white grapes, collected in the 2015 vintage in Titaguas (Spain) and stored in several small frozen volumes (4 l, −20*°*C). Before its use, the must was clarified by sedimentation for 24 h at 4*°*C and sterilized by adding dimethyl dicarbonate at 1 ml.l^−1^. All fermentations were performed in 3*x* independent biological replicates in 500 ml controlled bioreactors (MiniBio, Applikon, the Netherlands) filled with 470 ml of natural grape must. Each bioreactor was inoculated using an overnight starter culture cultivated in Erlenmeyer flasks containing 25 ml of YPD medium (2% glucose, 0.5% peptone, 0.5% yeast extract) at 25*°*C, 120 rpm in an agitated incubator (Selecta, Barcelona, Spain). Strain inoculation was done at OD600=0.100. The dynamics of the fermentation was registered using different probes and detectors to control and measure temperature, pH, dissolved oxygen (Applikon, The Netherlands) and effluent carbon dioxide level (INNOVA 1316 Multi-Gas Monitors, LumaSense Technologies). Data were integrated into the BioExpert software tools (Applikon, The Netherlands). The fermentation was complete when a constant sugar content was reached as measured by HPLC.

### 4.3 Sampling and quantification of extracellular metabolites

Extracellular metabolites, including sugars, organic acids, main fermentative by-products, and yeast assimilable nitrogen (YAN) were determined at ten sampling times during the fermentation. Residual sugars (glucose, fructose), organic acids (acetate, succinate, citrate, malate and tartrate) and the main fermentative by-products (ethanol, glycerol and 2.3 butanediol) were quantified using HPLC (Thermo Fisher Scientific, Waltham, MA) coupled with refraction index and UV/VIS (210 nm) detectors. Metabolites were separated through a HyperREZ XP Carbohydrate H+ 8 *μ*m column coupled with a HyperREZ XP Carbohydrate Guard (Thermo Fisher Scientific, Waltham, MA). The analysis conditions were: eluent, 1.5 mM of H_2_SO_4_; 0.6 ml.min^−1^ flux and a 50*°*C oven temperature. For sucrose determination, the same HPLC was equipped with a Hi-Plex Pb, 300 × 7.7 mm column (Agilent Technologies, CA, USA) and the following analysis conditions were used: eluent, Milli-Q water; 0.6 ml.min^−1^ flux and oven temperature of 50*°*C. The retention times of the eluted peaks were compared to those of commercial analytical standards (Sigma-Aldrich, Madrid, Spain). Metabolite concentrations were quantified by the calibration graphs (R2 value > 0.99) of the previously obtained standards from a linear curve fit of the peak areas using ten standard mixtures.

Determination of yeast assimilable nitrogen in the form of amino-acids and ammonia was carried out following the same protocol as Su et al. (2020). A volume of supernatant was removed from the fermenter, and amino acids and ammonia separated by UPLC (Dionex Ultimate 3000, Thermo Fisher Scientific, Waltham, MA) equipped with a Kinetex 2.6u C18 100A column (Phenomenex, Torrance, CA, USA) and Accucore C18 10 x 4.6 mm 2.6 *μ*m Defender guards (Thermo Fisher Scientific, Waltham, MA). For derivatization, 400 *μ*l of the sample was mixed with 430 *μ*l borate buffer (1M, pH 10.2), 300 *μ*l absolute methanol and 12 *μ*l of diethyl ethoxymethylenemalonate (DEEMM), and ultra-sonicated for 30 min at 20*°*C. The ultra-sonicated sample was incubated up at 80*°*C for 2 hours to allow the complete degradation of excess DEEMM. Once the derivatization finished, the sample was filtered with 0.22 *μ*m filter before injection. The target compounds in the sample were then identified and quantified according to the retention times, UV-vis spectral characteristics and calibration curves (R2 value > 0.99) of the derivatives of the corresponding standards. Amino acid standard (ref. AAS18), asparagine and glutamine purchased from Sigma-Aldrich were used for calibration.

### 4.4 Higher alcohols and esters

We also determined the concentrations of higher alcohols and esters for each sampling time. Volatile compound extraction and gas chromatography were performed following the protocol of Rojas et al. (2001). Extraction was performed using headspace solid phase-micro-extraction sampling (SPME) with polydimethylsiloxane (PDMS) fibers (Supelco, Sigma-Aldrich, Barcelona, Spain). Aroma compounds were separated by GC in a Thermo TRACE GC ULTRA chromatograph (Thermo Fisher Scientific, Waltham, MA) equipped with a flame ionization detector (FID), using a HP-INNOWAX 30 m x 0.25 mm capillary column coated with a 0.25 mm layer of cross-linked polyethylene glycol (Agilent Technologies, CA, USA). Helium was the carrier gas used (flow 1 ml.min^−1^). The oven temperature program was: 5 min at 60*°*C, 5*°*C.min^−1^ to 190*°*C, 20*°*C.min^−1^ to 250*°*C and 2 min at 250*°*C. The detector temperature was 280*°*C, and the injector temperature was 220*°*C under splitless conditions. The internal standard was 2-heptanone (0.05% w/v). Volatile compounds were identified by the retention time for reference compounds. The quantification of the volatile compounds was determined using the calibration graphs of the corresponding standard volatile compounds.

### 4.5 Physiological and biomass parameters

Physiological and biomass parameters, including OD600, dry weight (DW), colony-forming units (CFUs) and average cell diameter (ACD), were determined at each sample time, providing that the cell sample was sufficient to perform the corresponding measure. DW determination was performed by centrifuging 2 ml of the fresh sample placed in a pre-weighed Eppendorf tube in a MiniSpin centrifuge (Eppendorf, Spain) at maximum speed (13.200 rpm) for 3 min. After centrifugation, the supernatant was carefully removed, the pellet washed with 70% (v/V) ethanol, and centrifuged in the same conditions. After washing, the aqueous supernatant was removed carefully and the tube placed in a 65*°* C oven for 72h. DW was finally obtained by measuring the mass weight difference of the tube with a BP121S analytical balance (Sartorius, Goettingen, Germany). OD600 was measured at each sampling time using a diluted volume of sample and a Biophotometer spectrophotometer (Eppendorf, Germany). CFUs were determined using a 100-200 *μ*l of a diluted volume of samples plated in YPD solid medium (2% glucose, 2% agar, 0.5% peptone, 0.5%yeast extract) and incubated two days at 25*°* C. The resulting colonies were counted with a Comecta S.A Colony Counter. Only plates with CFUs between 30 and 300 were used to calculate the CFUs of the original sample. For ACD determination, a volume of cell sample was diluted into a phosphate-buffered saline solution and cell diameter measured using a Scepter Handled Automated Cell Counter equipped with a 40 *μ*m sensor (Millipore, Billerica, USA).

### 4.6 Orthology analysis and genome-scale metabolic reconstruction

Genomes of ScT73, SuBMV58 and SuCECT12600 were sequenced and assembled in previous works ((Morard et al., 2019); Macías et al., (unpublished)). Genome assemblies were annotated by homology and gene synteny using RATT (Otto et al., 2011). This approach let us transfer the systematic gene names of *S. cerevisiae* S288c annotation (Goffeau et al., 1996) to our assemblies and, therefore, to select only those syntenic orthologous genes in T73, CECT12600 and BMV58 genomes for subsequent analyses.

We added to the consensus genome-scale reconstruction of *Saccharomyces cerevisiae S288C (v.8.3.2)* metabolites and reactions related to amino acid degradation and higher-alcohols and esters formation. This refined model was then used as a template for reconstructing strain-specific genome-scale models for SuBMV58, SuCECT12600 and ScT73. MetaDraft, AuReMe and the results from the orthology analysis were used to create the strain-specific models.

### 4.7 Flux balance analysis

Flux balance analysis (FBA) (Varma and Palsson, 1994; Orth et al., 2010) is a modeling framework based on knowledge of reaction stoichiometry and mass/charge balances. The framework relies on the pseudo steady-state assumption (no intracellular accumulation of metabolites occurs). This is captured by the well known expression:

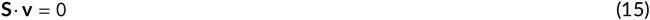

where **S** is stoichiometric matrix of (n metabolites by m reactions) and **v** is a vector of metabolic fluxes. The number of unknown fluxes is higher than the number of equations and thus the system is undetermined. Still it is possible to find a unique solution under the assumption that cell metabolism evolves to pursue a predetermined goal which is defined as the maximisation (or minimisation) of a certain objective function (*J*):

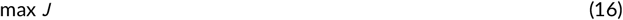

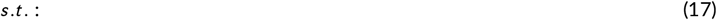

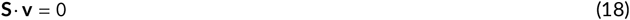

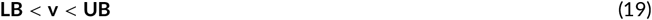

where **LB** and **UB** correspond to the lower and upper bounds on the estimated fluxes. Examples of objective functions *J* include growth rate, ATP, or the negative of nutrient consumption, etc.

Typically, multiple optimal solutions exist for a given FBA problem. In parsimonious FBA (pFBA), the result is the most parsimonious of optimal solutions, i.e., the solution that achieves the specific objective with the minimal use of gene products and the minimization of the total flux load (Machado and Herrgård, 2014).

### 4.8 Parameter estimation

The aim of parameter estimation is to compute the unknown parameters - growth related constants and kinetic parameters - that minimize some measure of the distance between the data and the model predictions. The maximum-likelihood principle yields an appropriate measure of such distance (Walter and Pronzato, 1997):

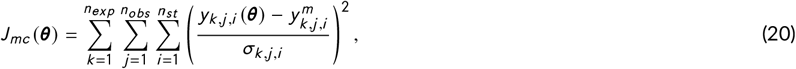

where *n*_*exp*_, *n*_*obs*_ and *n*_*st*_ are, respectively, the number of experiments, observables (measured quantities), and sampling times while *σ*_*k*,*j*,*i*_ represents the standard deviation of the measured data as obtained from the experimental replicates. 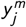 represents each of the measured quantities, *X* ^*m*^ and *C* ^*m*^ in our case, and *y*_*j*_ (***θ***) corresponds to model predicted values, *X* and *C*. Observation functions were included for *C F U s* and *OD* 600 in order to scale viable cell-mass (*X*_*v*_) and active cell-mass (*X*_*A*_), respectively.

Parameters are estimated by solving a nonlinear optimization problem where the aim is to find the unknown parameter values (***θ***) to minimize *J*_*mc*_ (***θ***), subject to the system dynamics - the model- and parameter bounds Balsa-Canto et al. (2010).

### 4.9 Uncertainty analysis

In practice, the value of the parameters ***θ*** compatible with noisy experimental data is not unique, i.e., parameters are affected by some uncertainty (Balsa-Canto et al., 2010). The consequence of significant parametric uncertainty is that it may impact the accuracy of model predictions.

To account for model uncertainty, we used an **ensemble approach**. To derive the ensemble, we apply the bootstrap smoothing technique, also known as bootstrap aggregation (the Bagging method) (Breiman, 1996; Bühlmann, 2012). The bagging method is a well established and effective ensemble model/model averaging device that reduces the variability of unstable estimators or classifiers (Büchlmann and Yu, 2002). The underlying idea is to consider a family of models with different parameter values **Θ** = [***θ***_1_ … ***θ***_*N*_]^*T*^ compatible with the data ***y***^*m*^, when using the model to predict untested experimental setups. The matrix of parameter values **Θ** consistent with the data is obtained using *N* realizations of the data obtained by bootstrap (Efron and Tibshirani, 1988). Each data realization has the same size as the complete data-set, but it is constructed by sampling uniformly from all replicates (3 biological replicates per sampling time). Within each iteration, each replicate has an approximate chance of 37% of being left out, while others might appear several times. The family of solutions, **Θ**, is then used to make *N* predictions (dynamic simulations) about a given experimental scenario. The median of the simulated trajectories regards the model prediction, while the distribution of the individual solutions at a given sampling time provides a measure of the uncertainty of the model.

### 4.10 Analysis of dynamic metabolic fluxes

We selected the most relevant metabolic pathways using a score, which provides a measure of the net flux over time during growth and stationary phases. In particular, we computed the integral of each flux multiplied by the biomass (mmol · h ^−1^) over time and normalised its value with the accumulated flux of consumed hexoses (glucose and fructose):

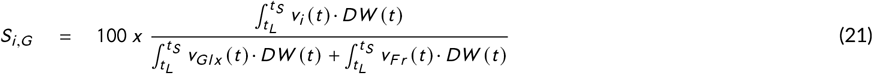

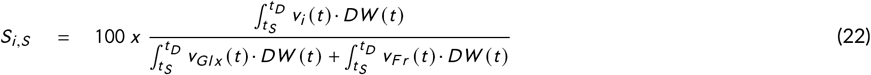

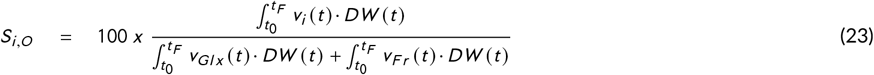

where *S*_*i*,*G*_ corresponds to the score of the flux *i* during growth, *S*_*i*,*S*_ corresponds to the score during the stationary, and *S*_*i*,*D*_ decay phases, *v*_*i*_ (*t*) (mmol · h ^−1^ · DW ^−1^) is the flux under scrutiny, *v*_*Gl x*_ (*t*) (mmol · h ^−1^ · DW ^−1^) is the flux of glucose, *v*_*F r*_ (*t*) (mmol · h ^−1^ · DW ^−1^) is the flux of fructose, and DW is the predicted dry-weight biomass (g). Results correspond to mmol of produced compound per mmol of consumed hexose x 100 (denoted as mmol/mmolH). Score values indicate the overall impact of each reaction in the net oxidation or reduction of electron carriers during the given phase of the fermentation.

### 4.11 Numerical tools

To automate the modeling pipeline we used the AMIGO2 toolbox (Balsa-Canto et al., 2016). To solve the dFBA problem we used a variable-step, variable-order Adams-Bashforth-Moulton method to solve the system of ordinary differential equations that describe the dynamics of the extracellular metabolites. At each time step the pFBA problem was solved using the COBRA Toolbox (Schellenberger et al., 2011). The global optimiser *Enhanced Scatter Search* (eSS, Egea et al. (2009)) was used to find the optimal parameter values in reasonable computational time.

The ensemble model generation procedure is computationally intensive. However, since each parameter estimation instance in the ensemble is an entirely independent task, we were able to solve this problem in less than a day using 60 CPU cores on a Linux cluster. These tasks were automated with the help of bash scripts and the Open Grid Scheduler. All the scripts necessary to reproduce the results are distributed (https://drive.google.com/drive/folders/102cMDQ6B9TZFRX0yIkPjivMrCdtEvQfd?usp=sharing).

## Author contributions

DH implemented the model. RM performed the experiments. SM performed the reconstruction of strain-specific genome-scale models. LGM assembled and annotated the genome. DH, SM and LGM participated in the gap-filling process of metabolic functions in the model and the development of the biomass equations. EBC, RPT, AQ, BT and EB coordinated the work and design of the study/pipeline. DH, RM, SM, LGM, EBC wrote the original manuscript. All authors read, edited, and approved the final paper.

## Expanded View Figure legends

**Expanded View Figure 1.**
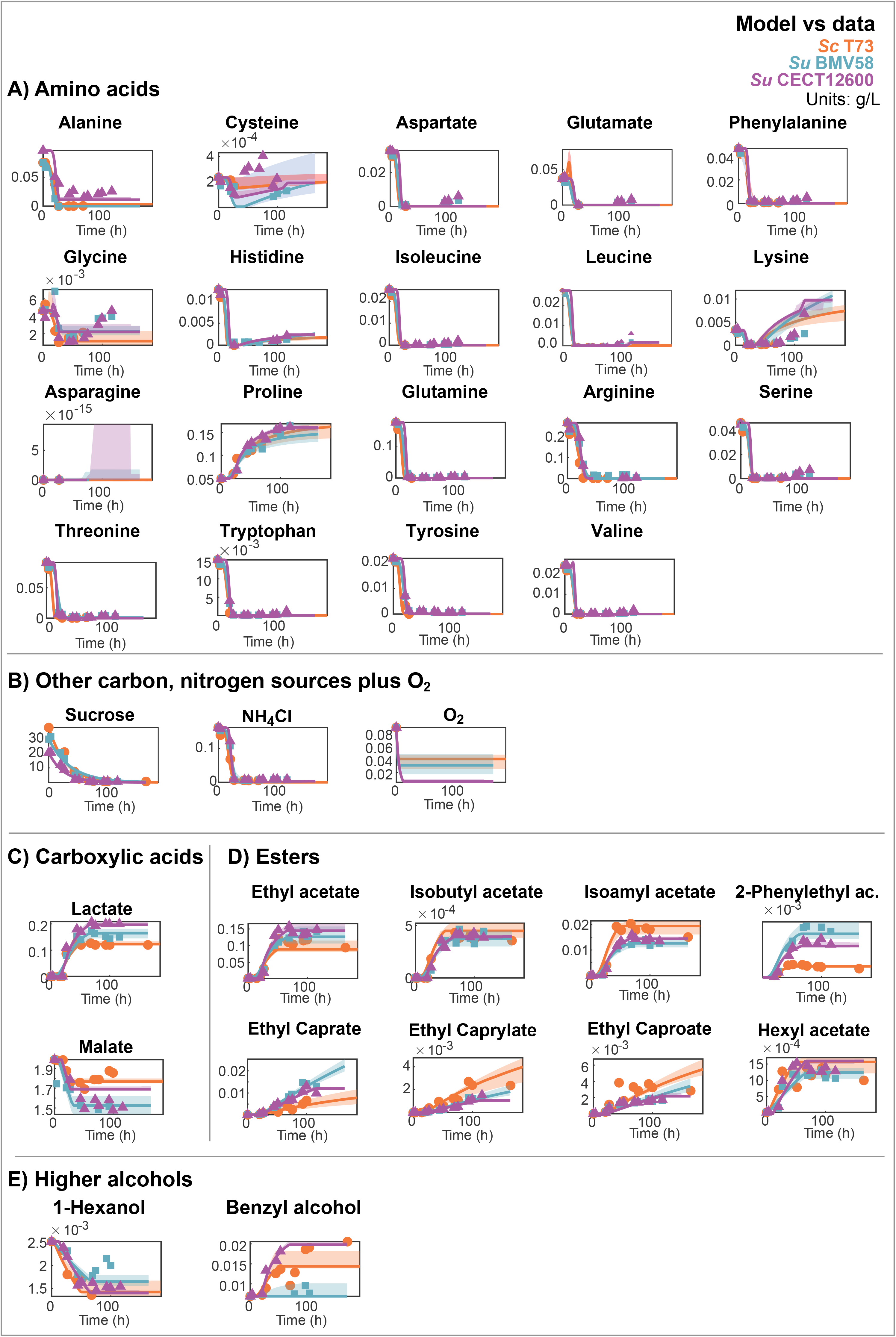
Model vs data for external metabolites: A) Amino acids, B) Other carbon, nitrogen sources plus O_2_, C) Carboxylic acids, D) Esters and E) Higher alcohols.

**Expanded View Figure 2.**
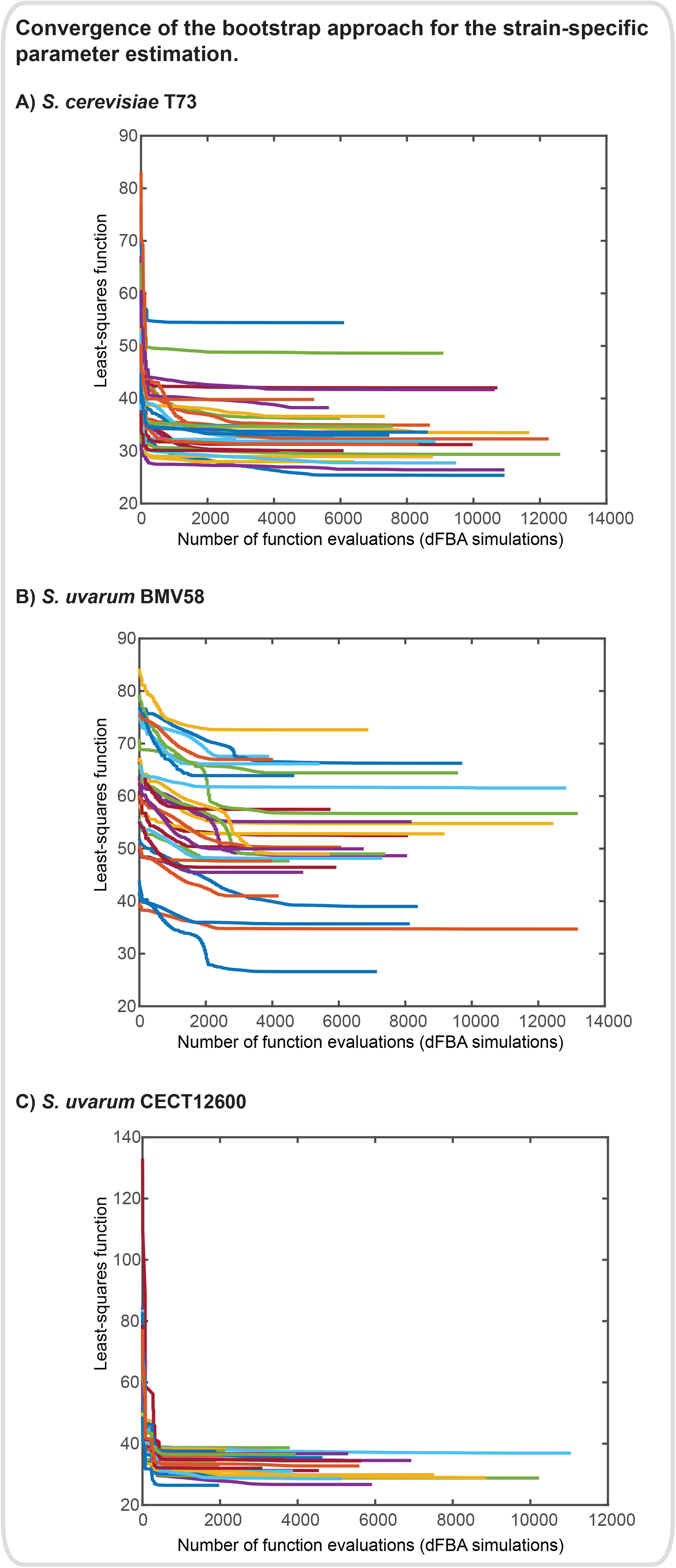
Convergence of the bootstrap approach for the strain-specific parameter estimation. Different bootstrap realizations result in different optimal least-squares values. Differences are more notorious for the case of SCT73 and SUBV58.

**Expanded View Figure 3.**
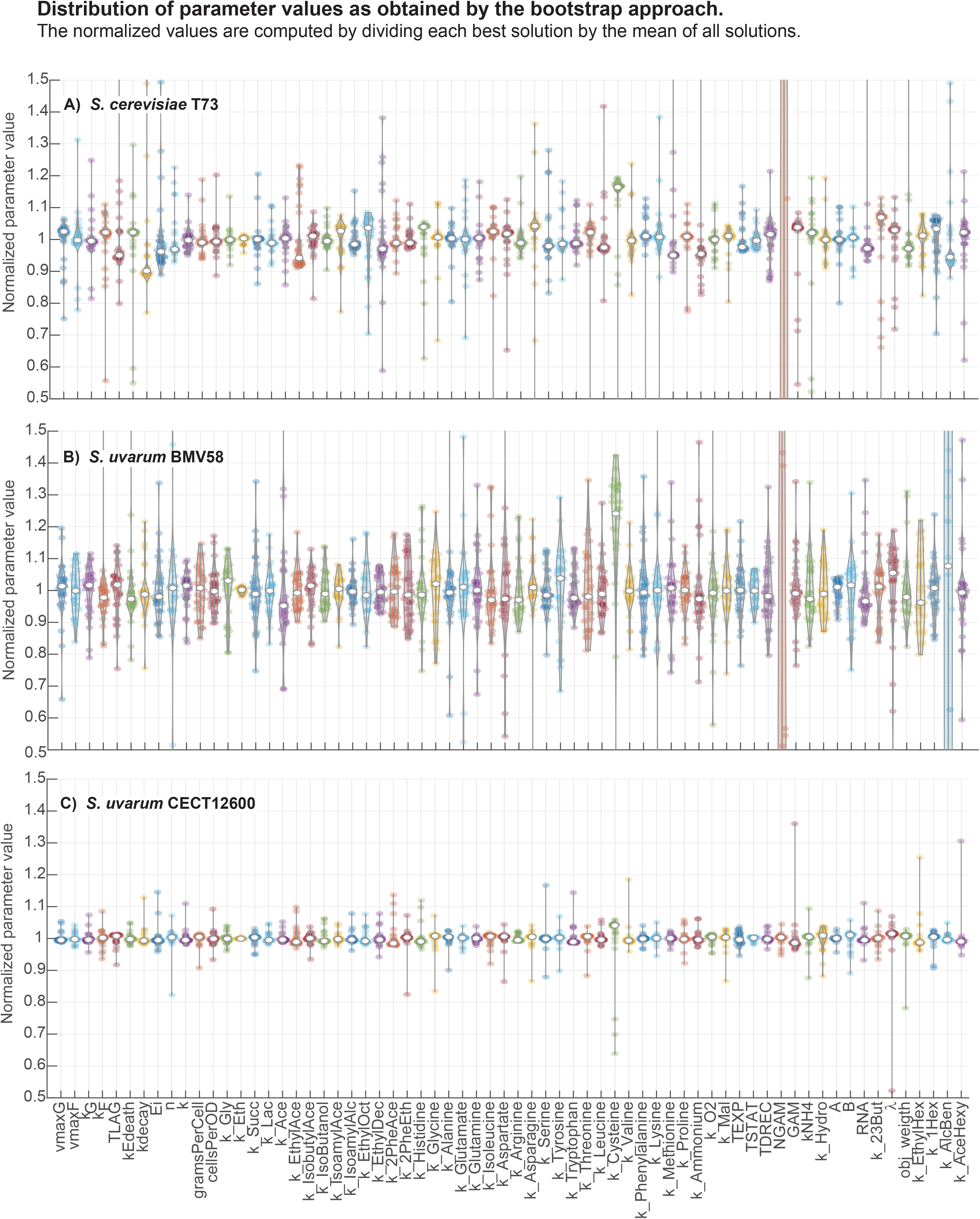
Distribution of the strain-specific parameter values as computed by the bootstrap approach. For most parameters, uncertainty is below the 20%. Remarkably this uncertainty does not translate into a high uncertainty on the model predictions.

**Expanded View Figure 4.**
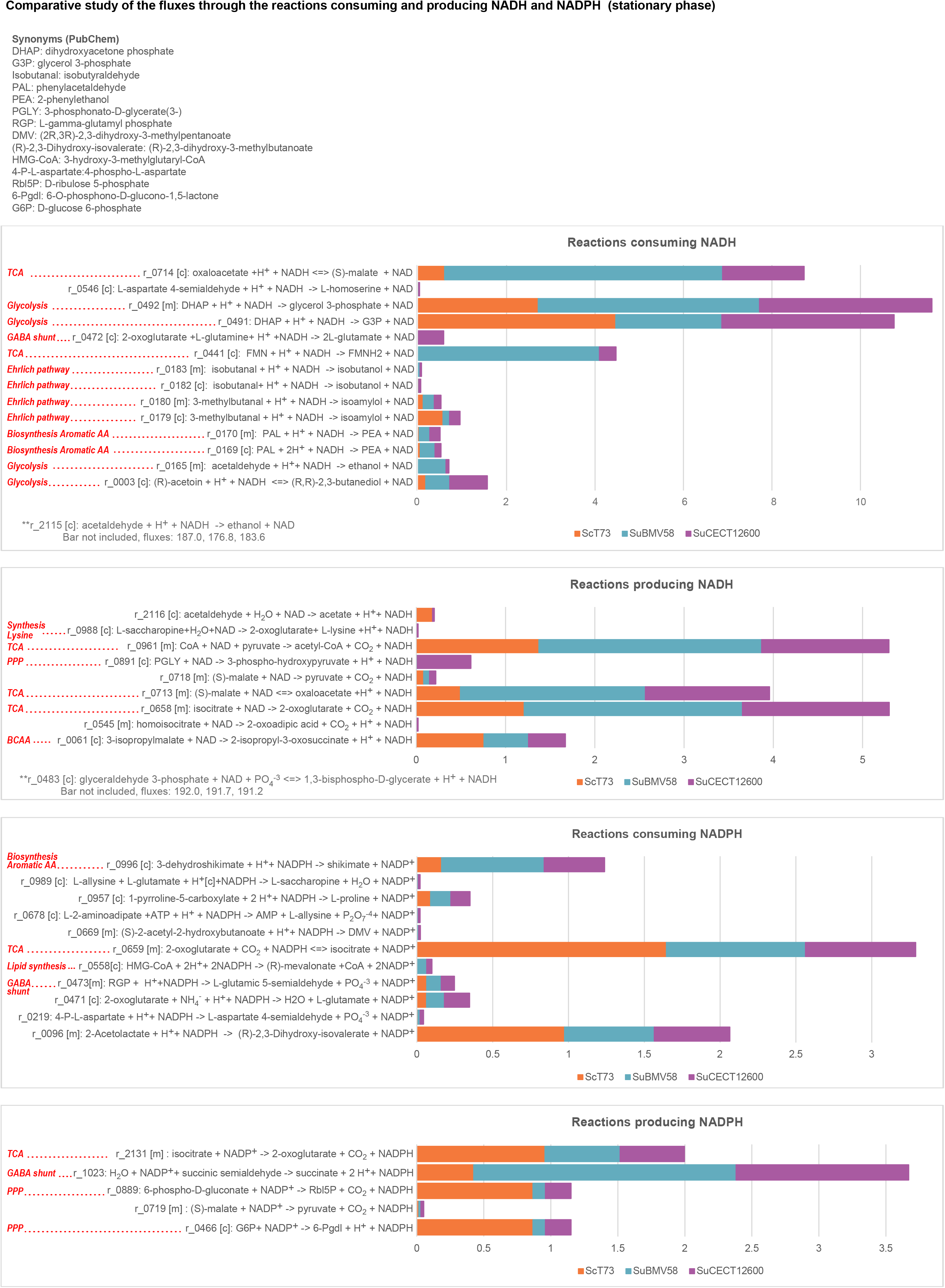
Comparative study of the fluxes through the reactions consuming and producing NADH and NADPH in the stationary phase for the three strains.

## Expanded View Table legends

- **Expanded View Table 1.** Reactions and metabolites added to the Yeast8 reconstruction.
- **Expanded View Table 2.** Descriptors of the aromas added to the Yeast8 consensus genome-scale reconstruction
- **Expanded View Table 3.** Multi-phase and multi-objective dynamic flux balance analysis. Definition of the objective function(s) and constraints per phase in the batch process.
- **Expanded View Table 4.** Model goodness-of-fit as measured by the R^2^ criterion for all measured states.
- **Expanded View Table 5.** Optimal parameter values as obtained by the bootstrap approach.
- **Expanded View Table 6.** Selected values of predicted fluxes for all batch process parts (growth, stationary, and decay) and overall fluxes. Values were selected considering the maximum of the values achieved for the three strains *max S*_*i*_ > 0.01 (with i, the number of flux).
- **Expanded View Table 7.** Fold-change of intracellular metabolites glutamate, GABA, GHB, succinate, fumarate and 2-oxoglutarate in synthetic wine fermentation. Data correspond to *S. cerevisiae* (QA23) grown at standard (28*°* C) and cold temperature (12*°* C) and *S. uvarum* at cold temperatures (12*°* C). Experimental procedures can be found on (López-Malo et al., 2013). Substantial accumulation of metabolites involved in the GABA shunt and PHB (i.e. GHB) synthesis was found in *S. uvarum* grown at low temperatures.

## APPENDIX

## Dynamic genome-scale modelling shows different redox balancing among yeast species in fermentation

## 1 Orthology analysis and genome-scale metabolic reconstruction

Genomes of ScT73, SuBMV58 and SuCECT12600 were sequenced and assembled in previous works [7], Macías et al., (unpublished)). Genome assemblies were annotated by homology and gene synteny using [8]. This approach let us transfer the systematic gene names of *S. cerevisiae* S288c annotation [4] to our assemblies and therefore, to select only those syntenic orthologous genes in T73, CECT12600 and BMV58 genomes for subsequent analyses.

We added to the consensus genome-scale reconstruction of *Saccharomyces cerevisae* S288C (v.8.3.1) metabolites and reactions related to amino acid degradation and higher-alcohols and esters formation. This refined model was then used as a template for reconstructing strain-specific genome-scale models for SuBMV58, SuCECT12600 and ScT73. First, AuReMe was used to generate draft genome-scale metabolic models for each strain using the refined model as a template. As a result, we obtained draft networks that included both gene-associated reactions (supported by genomic evidence and orthology) and non-gene associated reactions, such as transport reactions based on diffusion and exchange reactions, which were assumed to also occur in the strain-specific models. In addition, Metadraft was also used to generate draft networks for each strain using the refined yeast8 as a template. Reactions from MetaDraft were added to the drafts generated with AuReMe. Finally, the models were gap-filled using the refined template of yeast8 as the universe dataset from which reactions are taken to gap-fill the draft models. Figure S1 presents the differences between model reconstructions as compared to the consensus Yeast8.

**Figure S1:**
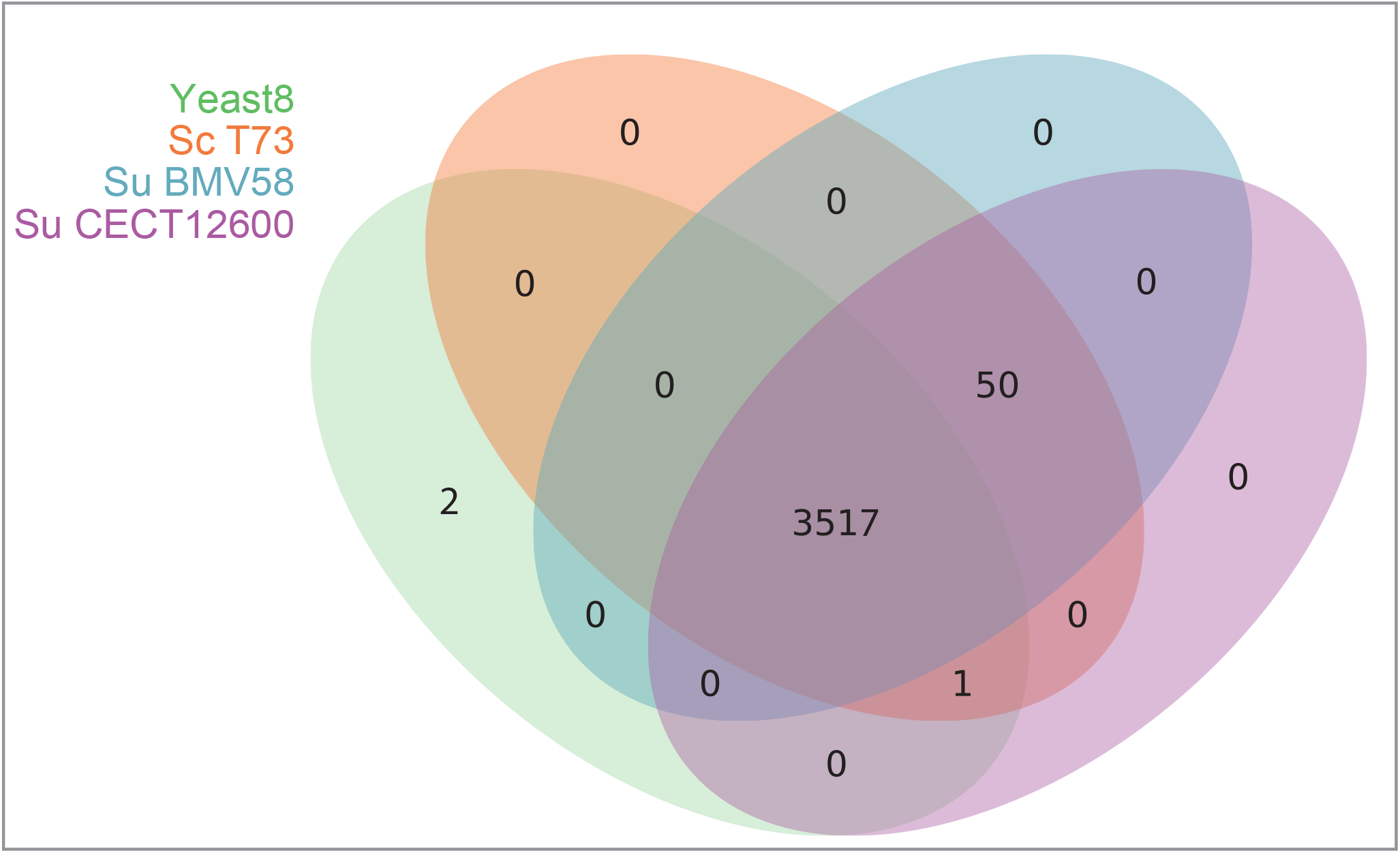
Differences between model reconstructions as compared to Yeast8.

## 2 Detailed description of the redox balance mechanisms used by the different strains

The comparative analysis of dynamic flux ratios showed that major differences between strains occur in the stationary phase. Our results show that *S. uvarum* and *S. cerevisiae* strains use different redox balance strategies markedly visible during the stationary phase of the fermentation. Here we present a detailed view of the differences found in the central carbon metabolism and the production of higher alcohols.

## 2.1 Central carbon metabolism

Figure S2 presents the dynamic flux ratios during the stationary phase of the most relevant pathways in the central carbon metabolism, including glycolysis and their impact on redox couple co-factors NADH/NAD+ and NADPH/NADP+.

## Production of ethanol

Taking the entire course of the fermentation in consideration, *S. uvarum* strains had a lower ethanol production rate than *S. cerevisiae* strain (187.40, 179.14, 182.40 *mmol/mmolH* in ScT73, SuBMV58 and SuCECT12600, respectively; Expanded view Table EV6, r 2115). Also, the model predicted slightly lower ethanol rates during growth phase for the three strains (173.10, 177.24, 183.34 *mmol/mmolH*), while they were quite similar during stationary phase (186.86, 177.24, 183.34 *mmol/mmolH*) (Figure S2.I; Figure EV5). Part of the ‘missing’ carbon was to be found in the yield of glycerol and in other downstream pathways using pyruvate as substrate.

## Production of glycerol

During the growth phase, the two *S. uvarum* strains had higher glycerol production rates, with SuBMV58 and SuCECT12600 strains producing 8.37 and 8.34 *mmol/mmolH* respectively, while the ScT73 strain only produced 7.25 *mmol/mmolH* (Table EV6, r 0489). Afterwards, the model predicted a smaller but still significant difference in glycerol production during the stationary phase (7.09, 7.84 and 7.81 *mmol/mmolH* for ScT73, SuBMV58 and SuCECT12600 respectively; Figure S2.I). Consistent with this, the overall score for the production of glycerol was lower for ScT73 (4.21 *mmol/mmolH*) than for both *S. uvarum* (> 6*mmol/mmolH*) (Table EV6, r 0489). As an NADH-consuming process, biosynthesis of glycerol plays an essential role in maintaining cytosolic redox balance during anaerobic conditions by oxidizing excess NADH to NAD+. Glycerol is also the only compatible solute to counterbalance the osmotic pressure in *S. cerevisiae* with glucose as a carbon source [1]. Directed evolution experiments exposing *S. cerevisiae* to osmotic stress led strains to produce a higher amount of glycerol, 2,3-butanediol and succinate [10]. In winemaking conditions, cells suffer hyperosmotic stress due to the elevated amount of sugars. The fact that *S. uvarum* strains present a higher amount of extracellular glycerol may indicate that *S. cerevisiae* tends to accumulate intracellular glycerol at early times during fermentation as previously reported by Perez-Torrado et al. [9].

## Production of succinate

The model predicted that the fraction of pyruvate incorporated into the mitochondria during the stationary phase accounted for 3.22, 3.64, 2.34 *mmol/mmolH* in ScT73, SuBMV58 and SuCECT12600 respectively (r 2034). Once inside mitochondria, pyruvate was either used for acetyl-CoA formation (1.37, 2.49, 1.43,*mmol/mmolH* for ScT73, SuBMV58 and SuCECT12600 respectively, r 0961) or directed towards the *de novo* production of valine and isobutanol (0.95, 0.59, 0.50, *mmol/mmolH* Figure S3.A).

Consistent with succinate raw data (Figure S2.E), the model estimated a significantly higher production rate of this by-product on the overall fermentative process in SuBMV58 (3.59 *mmol/mmolH*) and SuCET12600 (1.10 *mmol/mmolH*) than ScT73 (0.35 *mmol/mmolH*) (Table EV6, r 2057). Remarkably, the difference in production between *S. uvarum* strains was noteworthy. We also reported interesting variations in the rate and the origin of succinate according to the fermentation phase between strains. In anaerobic conditions during growth and stationary phases, there are three possible routes of producing succinate: the oxidative and reductive branch of the tricarboxylic acid cycle (TCA) GABA shunt. In anaerobic conditions, the TCA is truncated at the succinate level: TCA does not work cyclically but follows either a mitochondrial oxidative branch or a cytoplasmic reductive branch (Figure S2). In the former case, succinate is produced from pyruvate via four cytoplasmic reactions, yielding two NAD+ per pyruvate. In the other case, pyruvate is internalized into mitochondria and oxidized until succinate, producing three NADH per pyruvate. As for the GABA shunt, it is the pathway that circumvents the formation of succinyl-CoA in the oxidative branch of the TCA cycle. Instead, it converts 2-oxoglutarate into succinate with the intermediaries glutamate, GABA and succinate semi-aldehyde and yielding two NADH and one NADPH per pyruvate (Figure S2.I). In this regard, our model predicted that during the growth phase a carbon flux existed through the oxidative branch of TCA, but only until 2-oxoglutarate in the three strains and that there was no flux through the GABA shunt (Expanded View Table EV6). Therefore, the succinate formed during growth was yielded only by the reductive branch of TCA in ScT73, SuBMV58 and SuCECT12600 (Expanded View Table EV6). However, the most significant fraction of succinate was produced during the stationary phase, and with a significant higher rate by SuBMV58 (6.08 *mmol/mmolH*) and SuCECT1600 (1.66 *mmol/mmolH*) than ScT73 (0.43 *mmol/mmolH*). Interestingly, the model predicted that during stationary phase, the GABA pathway (0.42, 1.96 and 1.29 *mmol/mmolH* for ScT73, SuBMV58 and SuCECT12600 respectively; Figure S2.I; Figure EV4) summed to the reductive branch of TCA (0.00, 4.10 and 0.37 *mmol/mmolH* for ScT73, SuBMV58 and SuCECT12600 respectively; Figure S2.I; Figure EV4). Once again, the model suggested that during the stationary phase, the TCA oxidative branch was active until 2-oxoglutarate and the GABA shunt was responsible for completing the conversion of pyruvate-derived acetyl-CoA into succinate. Remarkably, the contribution of the TCA reductive branch succinate formation was almost twice than the GABA shunt in SuBMV58, while it was almost equal in ScT73 and SuCECT12600 strains. Also, as shown in Figure S2.I, the model suggested that the NADH produced in the oxidative branch of the TCA cycle could be re-oxidized at the level of the mitochondrial shuttles responsible for: i) the reduction of cytoplasmic DHAP to G3P (2.71, 5.00 and 3.90 *mmol/mmolH*), ii) the reduction of cytoplasmic acetaldehyde to ethanol (0, 0.63 and 0.07 *mmol/mmolH*).

**Figure S2:**
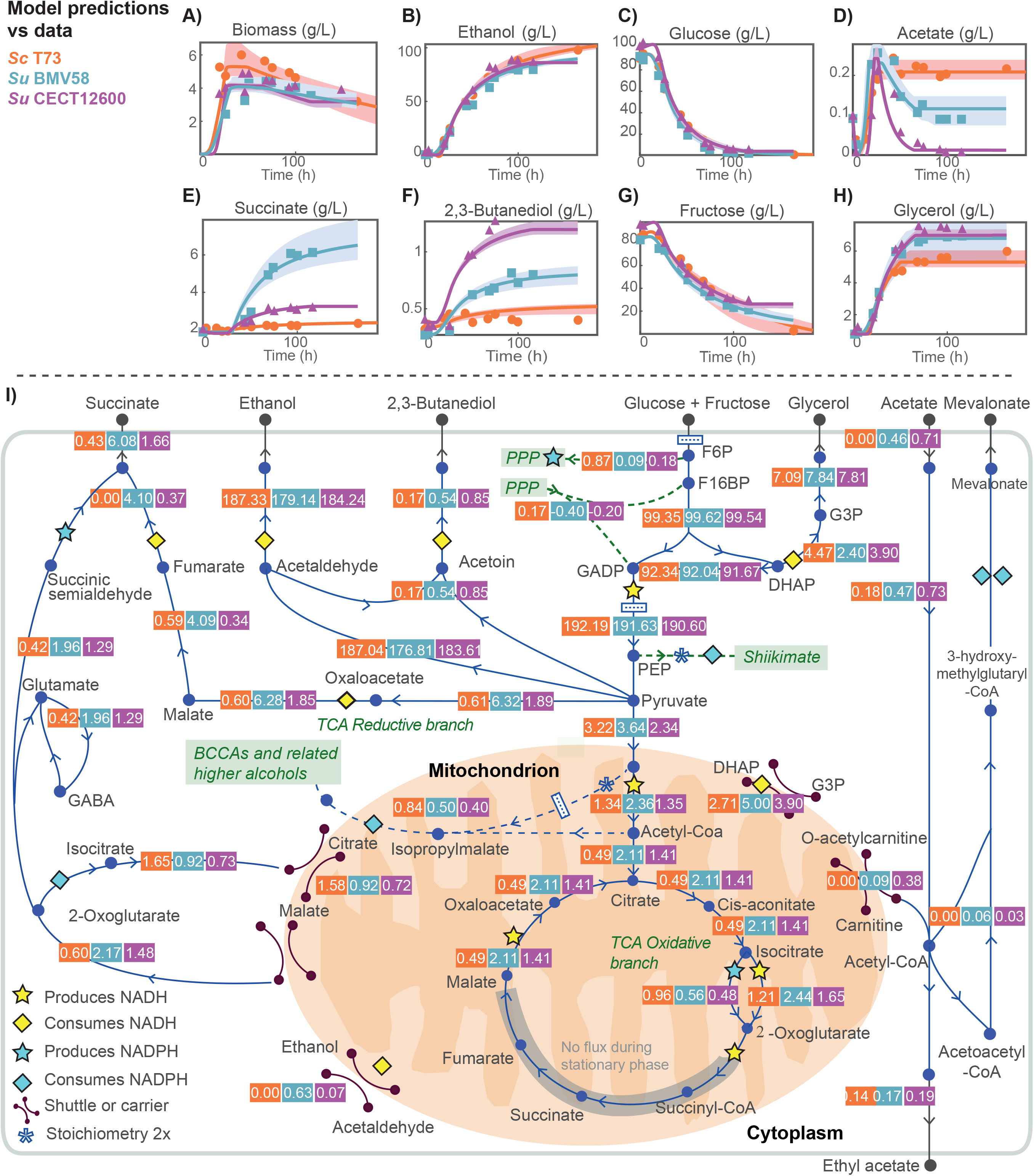
Redox balance in Central Carbon Metabolism. Figures A) to H) show model predictions versus the experimental data for biomass and extracellular metabolite concentrations associated with glycolysis and central carbon metabolism for the three strains. Figure I) shows the predicted intracellular dynamic flux ratios associated with the central carbon metabolism during the stationary phase. Figure I) illustrates how *S. uvarum* and *S. cerevisiae* strains use different redox balance strategies during the stationary phase of the fermentation. These differences result in the differential production of relevant external metabolites such as ethanol (B), succinate (E), 2,3-butanediol (F) or glycerol (H).

The finding that the GABA shunt would contribute to succinate formation in the case of *S. uvarum* strains was somewhat unexpected given previous results on *S. cerevisiae*. In fact, [2] concluded that for *S. cerevisiae* the GABA shunt was of little relevance on redox metabolism and that glutamate decarboxylase (GAD1) was poorly expressed when wine succinate is produced. Nevertheless, [5] observed substantial differences regarding intracellular levels of GABA in cryotolerant species. This fact and our results indicate that the production of succinate among *Saccharomyces* species requires further investigation.

## Production of 2,3-butanediol

*S. uvarum* strains were also more active in the production of 2,3-butanediol -a fermentative by-product involved in NADH oxidation from acetoin- than *S. cerevisiae* (Figure S2.I). The model correctly fitted extracellular raw data (Figure S2.F) and consistently predicted higher flux toward 2,3-butanediol production in *S. uvarum* strains during growth (0.22, 0.55, 0.98 *mmol/mmolH* in ScT73, SuBMV58 and SuCECT12600, respectively, r 1097) and stationary phases (0.17, 0.54 and 0.85 *mmol/mmolH* in ScT73, SuBMV58 and SuCECT12600, respectively). Again, but inversely to succinate, the flux difference towards 2,3-butanediol production between *S. uvarum* strains was noteworthy. Because both succinate and 2,3-butanediol generate from pyruvate (Figure S2.I), our results indicate that the two *S. uvarum* strains might use two different carbon redirection strategies around the pyruvate node: SuCECT12600 directing a larger fraction of pyruvate to the synthesis of 2,3-butanediol, and SuBMV58 to the synthesis of succinate.

## Production and consumption of acetate

Finally, we observed another striking difference between the three strains in the dynamics of acetate. As shown in Figure S2.D, the three strains produced acetate during the growth phase and until the entry into the stationary phase. Afterwards, while extracellular acetate concentration remained constant in ScT73, it decreased in both *S. uvarum* strains indicating an acetate consumption. Our model successfully described these phenotypes. On the first hand, we reported quite similar flux towards acetate production during the growth phase in the three strains (1.08, 1.14 and 1.42 *mmol/mmolH* in ScT73, SuBMV58 and SuCECT12600, respectively; Table EV6, r 1106). On the other hand, during the stationary phase, we obtained that SuBMV58 and SuCECT12600 strains consumed acetate with a rate of −0.46 and −0.71 *mmol/mmolH* respectively; on the contrary, ScT73 displayed an acetate rate equal to 0 *mmol/mmolH* (Table EV6, r 1106). Considering the entire fermentation process, we determined an overall acetate consumption of −0.12 *mmol/mmolH* for SuCECT12600, while a limited production of 0.036 *mmol/mmolH* was computed for SuBMV58. In the case of ScT73 strain, this production was five times higher (0.17 *mmol/mmolH*; Table EV6,r 1106).

In our first simulations, the model determined that *S. uvarum* strains used a significant part of this consumed acetate to produce succinate through the glyoxylate pathway. This result made sense from a pFBA point of view because this path resulted in the smallest overall flux throughout the metabolic network. However, we could not find literature supporting the production of glyoxylate in the presence of a large glucose concentration. Therefore, we decided to constrain the isocitrate lyase flux to zero. The revised model then suggested that during stationary phase SuBMV58 and SuCECT12600 strains incorporated the acetate derivative, acetyl-CoA, into ethyl acetate in the cytoplasm (0.17 and 0.19 *mmol/mmolH*); shifted most of the remaining fraction (0.09 and 0.38 *mmol/mmolH*) into the mitochondria through the carnitine shuttle system and the last part (0.06 and 0.03) was directed towards mevalonate (Figure S2.I). The fact that an amount of the acetate carbon was directed towards mevalonate is in line with recent experimental work by [6].

Inside the mitochondria, acetyl-CoA was used to form isopropylmalate (precursor of isoamyl alcohol, Figure S3.A) or further used in the TCA towards the synthesis of 2-oxoglutarate (Figure S2.I).

## Contributions to redox balance

All the aforementioned fermentative by-products (ethanol, glycerol, succinate, 2,3-butanediol and acetate) impact NADH/NAD+ and/or NADPH/NADP+ balance (Figure S2.I). However, their relative contribution to maintaining co-factors equilibrium varies according to the stoichiometry of the reaction, the function of the cell compartment in which they are produced (cytoplasm or mitochondria) and the activity of redox shuttles between compartments. Most of the glycolytic pyruvate was directed towards ethanol production for the three strains, known to be redox neutral. However, we noticed that both glycerol synthesis and reductive succinate production were more pronounced in *S. uvarum* strains during the stationary phase. Thus, according to the stoichiometry and localization of these pathways, it may result in a cytoplasmic surplus of NAD+ that should be compensated elsewhere in the metabolism of *S. uvarum* strains.

## 2.2 Production of higher alcohols

Higher alcohol production started during the growth phase and ceased at the end of the stationary phase. Our results reflect substantial differences in the accumulation of some higher alcohols between strains. In particular, the model predicted that the carbon skeletons of isoamyl alcohol, isobutanol and 2-phenyl ethanol were in a significant part synthesized *de novo* from glycolytic and pentose phosphate pathway intermediates, rather than coming from the catabolism of precursor exogenous amino acids (leucine, valine and phenylalanine respectively). Furthermore, the model shows that isoamyl alcohol and 2-phenyl ethanol contribute to glycerol formation in wine fermentation.

Figure S3.A shows the predicted intracellular fluxes related to higher alcohols during the stationary phase and its corresponding impact on the redox co-factors balance NADPH/NADP+ and NADH/NAD+. Readers can find the dynamic flux ratios in the Expanded view Table EV6. Figures 3.B-3.D correspond to the comparison between model predictions and raw measures of higher alcohols.

During the stationary phase, *S. uvarum* strains produced more 2-phenylethanol than ScT73 strain per unit of consumed hexoses (0.16, 0.66 and 0.38 *mmol/mmolH* for ScT73, SuBMV58 and SuCECT12600 respectively, r 1590) while the opposite pattern was observed for isoamyl alcohol (0.74, 0.48 and 0.39 *mmol/mmolH* for ScT73, SuBMV58 and SuCECT12600 respectively, r 1863). In contrast, the model prediction was quite similar for the three strains for isobutanol (~ 0.09 *mmol/mmolH*), as well as for other higher alcohols such as methionol and tyrosol which seemed to accumulate in minimal quantities (~ 0.001 *mmol/mmolH*) in response to perturbations in the amino acid pool.

## Production of 2-phenylethanol

*De novo* synthesis of 2-phenylethanol (PEA, Figures S3.A, S3.B) from carbohydrates contributed to redox homeostasis. Erythrose-4-phosphate (E4P, Figure S3.A) and phosphoenolpyruvate (PEP) are two sugar-phosphate intermediates of the pentose phosphate pathway (PPP) and glycolysis. They are the starting substrates of the chorismate pathway that lead to phenylalanine, which can subsequently be catabolized to 2-phenylethanol (Ehrlich pathway). From the beginning of the chorismate pathway to 2-phenylethanol, one NADPH (3-dihydroshikimate → shikimate) and one NADH (phenylacetaldehyde (PAL) → PEA) are consumed (Figure S3.A).

However, to fully understand the impact of *de novo* production of 2-phenylethanol on redox homeostasis, it is also important to remember that the PPP consists of two branches: i) an oxidative and irreversible branch from glucose-6-phosphate (G6P) to ribulose-5-phosphate (Rbl5P) resulting in the net formation of two reduced NADPH co-factors per molecule of glucose-6-phosphate (Figure S3.A), and ii) a non-oxidative branch consisting of reversible carbon shuffling reactions between sugar-phosphate molecules leading to important precursor metabolites (e.g. ribose-5-phosphate (R5P) and eryhtrose-4-phosphate (E4P)) and glycolytic intermediates (e.g. fructose-6-phosphate (F6P) and glyceraldehyde-3-phosphate (G3P)). Thus, the transketolase and transaldolase enzymes of this branch of the PPP provide a reversible link between the PPP and glycolysis [3].

**Figure S3:**
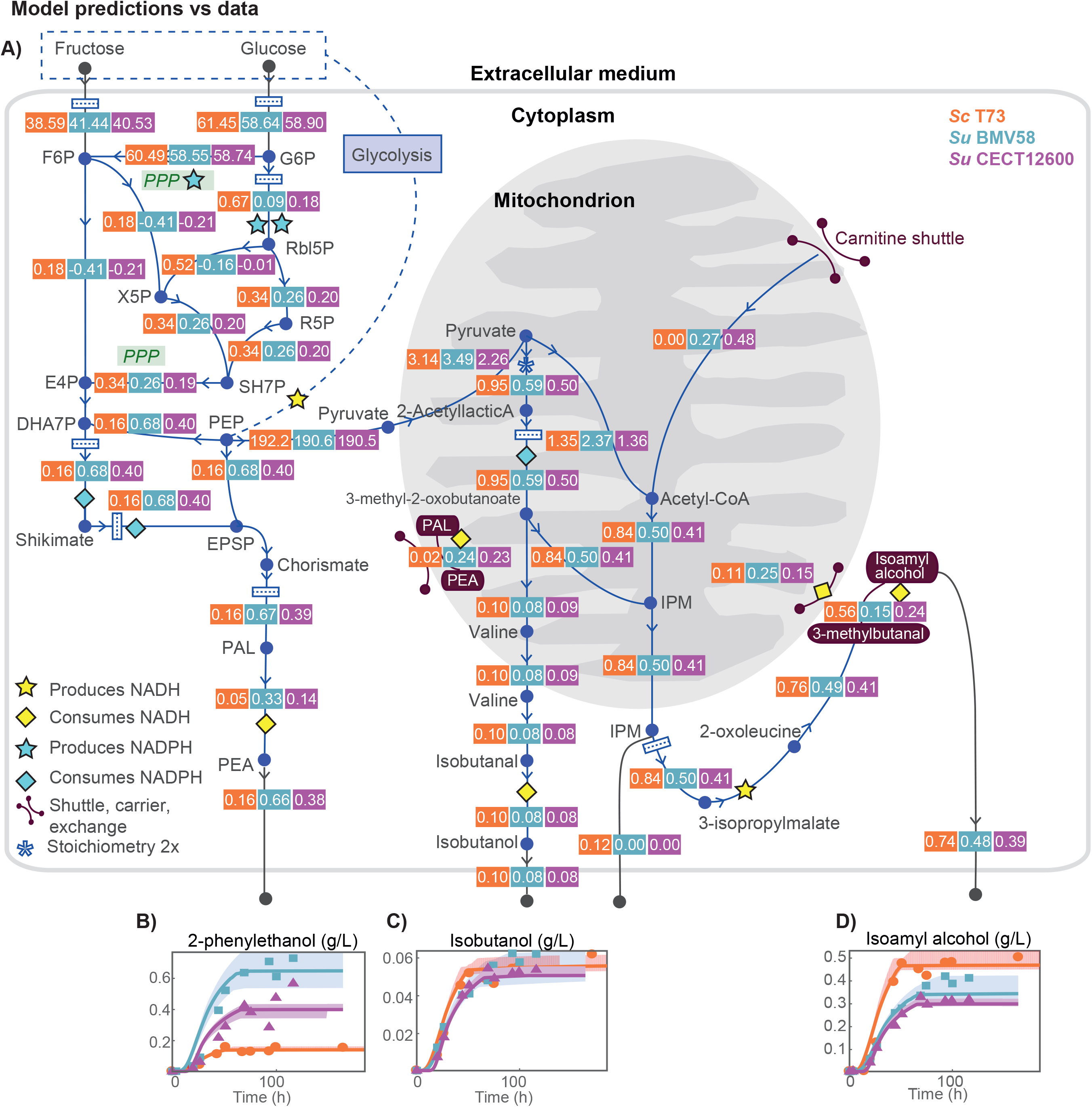
Redox balance in the production of higher alcohols. Figure A) shows the predicted intracellular fluxes related to higher alcohols during the stationary phase and its corresponding impact on the redox co-factors balance NADPH/NADP+ and NADH/NAD+. Figures 3.B-3.D correspond to the comparison between model predictions and raw measures of 2-phenylethanol, isobutanol and isoamyl alcohol.

In this context, if E4P required for *de novo* synthesis of PEA was generated trough the oxidative branch of PPP, PEA production would result in the accumulation of NADPH. However, if E4P was generated by the non-oxidative branch of PPP from sugar phosphate intermediates of the glycolysis, we would expect an NADP+ accumulation.

As shown in Figure S3.A, our model predicted that during stationary phase ScT73 had a greater flux through the oxidative branch of the PPP than *S. uvarum* strains (0.67, 0.09 and 0.18 *mmol/mmolH* for ScT73, SuBMV58 and SuCECT12600 respectively). On the contrary, dynamic flux ratios through several non-oxidative reactions of the PPP were higher in both *S. uvarum* strains (F6P → E4P : 0.18, −0.41 and −0.21 *mmol/mmolH*; F6P → X5P :0.18, −0.41 and −0.21 *mmol/mmolH*; Figure S3.A). Consistent with an higher 2-phenylethanol synthesis by SuBMV58 and SuCECT12600, the model predicted an increase in flux towards chorismate synthesis in both *S. uvarum* strains (0.17, 0.68 and 0.40 *mmol/mmolH* for ScT73, SuBMV58 and SuCECT12600, respectively). The higher flux through the PPP oxidative branch in ScT73 can be partially explained by NADPH requirement (at the level of 3-methyl-2-oxobutanoate formation), in the *de novo* synthesis of isoamyl alcohol. On the other hand, the alternative non-oxidative PPP strategy used by *S. uvarum* strains may contribute to providing NADP^+^ co-factors required in the NADP^+^-dependent glutamate degradation to succinate in the GABA shunt (0.42, 1.96 and 1.29 *mmol/mmolH* for ScT73, SuBMV58 and SuCECT12600, respectively; Figure S2.I).

## Production of isoamyl alcohol

During the stationary phase, the model predicted a substantial flux from pyruvate to 2-acetyllactic acid inside the mitochondria (0.95, 0.59 and 0.50 *mmol/mmolH* for ScT73, SuBMV58 and SuCECT12600 respectively; Figure S3.A; r 0097). In this reaction, two pyruvates are required per 2-acetyllactic molecule formed. The model also suggested that 2-acetyllactic acid was mainly directed to forming 3-isopropylmalate (IPM, 0.84, 0.50 and 0.41 mmol/mmolH for ScT73, SuBMV58 and SuCECT12600 respectively; Figure S3.A; r 0025), consuming one NADPH and one Acetyl-CoA. 3-isopropylmalate was mainly converted in isoamyl alcohol (0.76, 0.49 and 0.41 mmol/mmolH for ScT73, SuBMV58 and SuCECT12600 respectively) rather than for *de novo* leucine synthesis. The conversion of 3-isopropylmalate into 2-oxoleucine releases one NADH (r 0061), consumed during the reduction of 3-methylbutanal to isoamyl alcohol (Figure S3.A, r 0179). Following these reactions, the formation of each isoamyl alcohol molecule consumes two pyruvate molecules, one acetyl-CoA and one NADPH (Figure 3.A).

Besides, the formation of two pyruvates from one glucose release two NADH, and two NADH are produced per acetyl-CoA formed. Summing up, *de novo* synthesis of one isoamyl alcohol should result in excess of four NADH and one NADP^+^. Accordingly, this *de novo* synthesis of isoamyl alcohol from pyruvate has a relevant impact on redox balance. The citrate/2-oxoglutarate could provide the NADP^+^ required to synthesize 3-methyl-2-oxobutanoate inside the mitochondrion NADPH shuttle.

Remarkably, the production of higher alcohols contributed substantially to redox metabolism related to glycerol accumulation. During stationary phase, approximately 43% of the glycerol produced by the ScT73 strain was attributable to NADH derived from isoamyl-alcohol and 2-phenyl ethanol production. In the cases of SuBMV58 and SuCECT12600 strains, these values dropped to 36% and 27%, respectively.

